# Neural cell injury pathology due to high-rate mechanical loading

**DOI:** 10.1101/2021.05.12.443823

**Authors:** Jonathan B. Estrada, Harry C. Cramer, Mark T. Scimone, Selda Buyukozturk, Christian Franck

**Author notes:** These authors contributed equally to this work (Jonathan B. Estrada, Harry C. Cramer).

## Abstract

Successful detection and prevention of brain injuries relies on the quantitative identification of cellular injury thresholds associated with the underlying pathology. Here, by combining a recently developed inertial microcavitation rheology technique with a 3D *in vitro* neural tissue model, we quantify and resolve the structural pathology and critical injury strain thresholds of neural cells occurring at high loading rates such as encountered in blast, cavitation or directed energy exposures. We find that neuronal dendritic spines characterized by MAP2 displayed the lowest physical failure strain at 7.3%, whereas microtubules and filamentous actin were able to tolerate appreciably higher strains (14%) prior to injury. Interestingly, while these critical injury thresholds were similar to previous literature values reported for moderate and lower strain rates (< 100 1/s), the pathology of primary injury reported here was distinctly different by being purely physical in nature as compared to biochemical activation during apoptosis or necrosis.

**Teaser:** Controlled microcavitation enables quantitative identification of injury thresholds in neural cells.

## Introduction

Mechanical forces play a critical role in maintaining the form and function of many biological systems from single cells to entire organisms [1, 2]. From the perspective of a cell, maintenance of physical homeostasis involves a constant balancing act between internal and external forces and deformations. Critical cellular processes such as tissue morphogenesis [3], protein expression [4], and cell differentiation [5], are often a result of slight and well-coordinated changes in a cell’s physical homeostasis. In cases of significant disruption in homeostasis, such as in physical trauma, cells can produce a spectrum of insult-dependent pathologies, broadly classifiable as either immediate mechanical breaking of cellular microstructural components (primary injury), or the activation of apoptotic or necrotic pathways (secondary injury) [6, 7].

Abnormal loading conditions can occur from unintentional causes (e.g., during Traumatic Brain Injuries or TBI), or through intentionally administered therapeutic treatments including many modern surgical or disease-mitigating procedures [8]. Particularly, the last decade has seen rapid growth in the use of focused energy techniques including diagnostic [9–11] and therapeutic ultrasound [12–15] and laser-based ablation methods [16, 17] for a variety of patient treatment procedures, from removing tumors to correcting eyesight. The physical mechanism of tissue removal in ultrasound-based procedures such as histotripsy is the generation of cavitation bubbles that impart extremely high-rate and large deformations onto the surrounding cells and tissues which lead to tissue fragmentation, lysis and cell death. Laser-based ablation procedures can also produce significant microcavitation while heating the tissue [16–18]. Despite these methods’ growing popularity and the well-defined qualitative understanding of their use, little is quantitatively known about the relationship between bubble size, bubble pressure, and resulting tissue stresses and strains on cellular and tissue damage and injury [19].

Recently, blast-related traumatic brain injuries in the armed forces have pointed toward the possibility of a brain injury signature distinctively different from those associated with the pathologies of commonly diagnosed blunt-force mild to severe TBIs [20]. The chief distinction between these types of TBI could be the significantly higher deformation or strain rates experienced under such conditions, similar to those occurring during exposure to either electromagnetic or sonic-based directed energy devices [21]. To this end, a variety of efforts aimed toward mitigating or preventing TBI have turned to advanced computational models of the head and neck to better understand the role of high-rate deformations, forces, stresses and strains in the overall associated injury risk potential [22,23]. While these models have progressed in their complexity and predictive power, their experimental validation—specifically resolution of critical cellular and subcellular injury or damage thresholds—has remained elusive.

Considering that mechanical loading on the tissue and cells can be highly disruptive and destructive, it is paramount to resolve critical thresholds in force and deformation along with cellular pathologies under such conditions. Understanding the extent of disruption and injury is especially important in neural tissues to prevent any loss of function within the central nervous system.

Careful resolution and quantification of the mechanical forces and deformations, or strains, that lead to injurious cellular disruptions and possible cell death have remained a formidable challenge in practice. This is in part due to the fast, high-rate character of the mechanical deformations imparted onto the cells, which require camera frame rates on the order of hundreds of thousands to millions of frames per second to resolve. Furthermore, in order to remove loading and boundary artifacts that could complicate the post analysis of cellular injury, sample preparation and mounting cannot be injurious to the cells and tissues. This poses a major challenge to most traditional mechanical loading stages or bioreactors. Finally, and especially for neural tissues, scaffolds or tissue structures promoting three-dimensional (3D) cellular architectures are crucial for replicating physiologically relevant mimics of the *in vivo* tissue microenvironment.

To address these key challenges and to provide a robust method for quantitatively resolving cellular injury thresholds and pathologies with high spatial resolution, we combine our recently developed inertial microcavitation rheology (IMR) technique [24] with an established 3D *in vitro* neural cell tissue model [7]. Through 3D-positioning of a spatially-controlled inertial cavitation bubble within the 3D neural tissue model, high-rate mechanical deformations (material strain rates on the order of 10^3^ − 10^8^ 1/s), are achieved without the need to handle, mount or condition the tissue prior to loading. By integrating this technique into the optical path of a multiphoton microscope, high-resolution, 3D volumetric image stacks can be acquired before and after an induced high-rate loading event. To construct a strain- and strain-rate-dependent pathology atlas, common immunocytochemical markers are employed and correlated to the applied high-rate deformations. Additionally, by employing our previously developed quantitative theoretical framework built into IMR, we can estimate cellular strains and provide stress threshold estimates for cellular projections containing each subcellular structural component represented in the injury pathology. Taken together, this study provides a significant quantitative advancement in our understanding of this biologically and clinically important injury pathology, and provides actual, measured cell-level details on the critical strain and stress thresholds associated with structural degeneration across key cellular constituents including microtubules, filamentous actin, and microtubule-associated protein-2 (MAP2).

## Results

To resolve the pathology and critical injury thresholds of high-rate deformations on neural cell populations, a 3D *in vitro* neural tissue model [7] was integrated with our recently developed, laser-based microcavitation technique as a means to mechanically load the cellular constructs. The *in vitro* model consists of neural cells dissociated from the cortices of postnatal p0-p1 Sprague Dawley rat pups and encapsulated in a collagen-I hydrogel in a glass-bottomed 48 well plate. The neural cell population undergoes neurogenesis over the course of 7 days *in vitro* during which the cells form healthy network connections consistent with previous work [7, 25] as seen in Fig. 1a. Cells were determined to be well-encapsulated by collagen fibers using reflectance confocal and auto-fluorescence microscopy (Fig. 1b). To mechanically deform the cells at high loading rate, a single microcavitation bubble was generated inside the neural tissue model via a spot-focused, ~4 nanosecond pulsed laser aligned into the back port of an inverted microscope. Briefly, the deposited laser energy nucleates a microcavitation bubble, which grows and rapidly stretches the surrounding neural tissue as it expands to a maximum radius, *R*_max_ (Fig. 1c). The general bubble dynamics feature several, exponentially decaying oscillatory expansion-collapse cycles [24] until the bubble reaches its final stress-free, quasi-static equilibrium bubble radius, *R*_0_, over the course of ~ 120*μ*s. High-speed video of bubble dynamics from each experiment was recorded by one of two methods: at 270,000 frames per second using a Phantom v2511 (Vision Research Inc, Wayne, NJ) high-speed camera, or at 1 million frames per second with a Shimadzu HPV-X2 ultra-high speed camera (Shimadzu, Kyoto, Japan), both using bright field microscopy. An example set of high-speed images at cropped spatial and temporal resolution from the latter is shown in Fig. 1c. Bubble radii were determined from these images using a custom image processing algorithm modified from our prior work [24].

**Figure 1:**
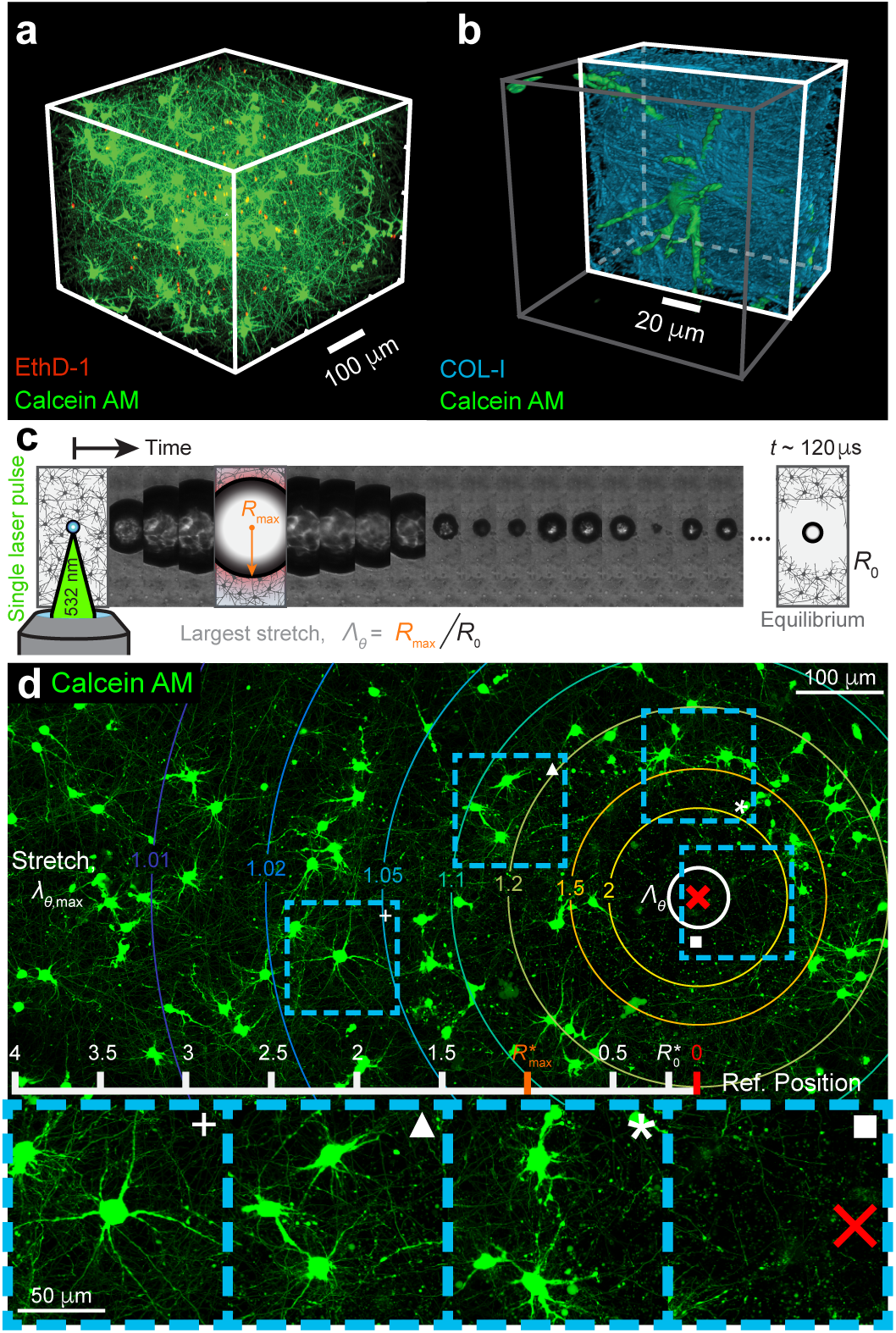
Experimental injury platform and projected, 3D micrographs of high loading rate cellular pathology. (a) 3D type I collagen gels are fabricated *in vitro*, containing primary neural cells (green) which grow and form healthy, connected networks. (b) Cells (green) are well-encapsulated by type I collagen (sky blue). (c) A single, 4 ns laser pulse focused through a microscope objective induces microcavitation in the gel. The cavitation microbubble rapidly grows in the sample, reaching a maximum radius *R*_max_ (orange tick mark), and subsequently collapses and oscillates, finally reaching a quasi-equilibrium state *R*_0_. (d) Cells in the vicinity of the bubble are rapidly deformed by radially-decaying circumferential (hoop) stretch levels, *λ_θ,_*_max_, indicated by the color-coded radial rings. Each radial ring represents the maximum tensile (hoop) stretch, *λ_θ_* = *λ_θ,_*_max_ experienced by each cell. While cells that experience stretches less than *λ_θ,_*_max_ < 1.05 remain morphologically intact (plus), cells closer to the bubble center exhibit varying degrees of degenerate projections 1.05 < *λ_θ,_*_max_ < 1.2 (triangle) and somatic disruption 1.2 < *λ_θ,_*_max_ < 2 (star, inset image rotated). Very near to the bubble epicenter (red x), cell somas and projections are entirely fragmented at *λ_θ,_*_max_ > 2, (square, inset image rotated). The overall spatial extent of injury is dependent on the maximum bubble size (*R*_max_), and is generally contained within 2–2.5×*R*_max_ as indicated by the overlaid ruler (units are in multiples of *R*_max_).

To quantify the physical deformations that the neural cells experience during high-rate cavitation loading we begin by describing the change in position, or displacement, of each material point (cells and collagen) using a standard continuum mechanics representation [24]. We denote the current radial position of the bubble wall in time as *R*(*t*), with an associated “reference” radius *R*_0_ of the bubble in the stress-free, equilibrium configuration at long times. The maximum radius, *R*_max_ is then defined as, *R*(*t* = 0) = *R*_max_. Radial positions in the material are then functions of both the bubble radius and time, and are denoted as *r*(*r*_0_*, t*) and *r*_0_ in the current and reference configurations, respectively. To relate the physical stretch experienced by the cells to the measured bubble radii *R*(*t*) during cavitation, we first define the material deformation gradient tensor **F**, as

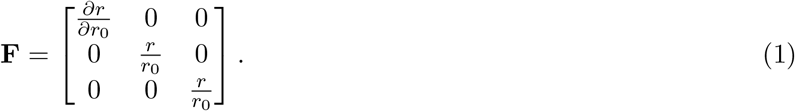

Considering the high-rate of loading, the material can be treated to be near-incompressible [26,27], i.e., det(**F**) = 1, yielding the following relationship:

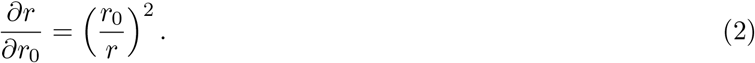

Equation 2 can then be solved in terms of the current radial position *r*(*r*_0_*, t*) as

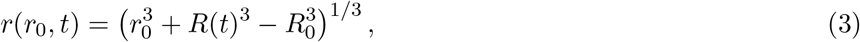

establishing a geometric relationship between any spatial position within the gel, *r*, and the current and equilibrium bubble radii, *R* and *R*_0_, respectively. Finally, we define the circumferential, or hoop, material stretch, *λ_θ_*, during mechanical loading as

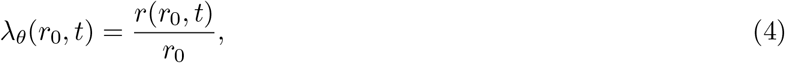

with the average maximum tissue stretch occurring at the bubble wall over all experiments during the time of largest bubble expansion defined as Λ_*θ*_,

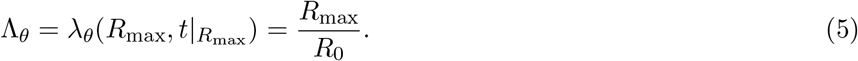

While a similar relationship can be established for the radial stretches, we focus on the predominantly tensile hoop (or circumferential) stretch, which prior work established to be one of the most injurious modes of deformation to cells [6]. Figure 1d provides representative projected micrographs of neural cells labeled with the live cell indicator Calcein AM that underwent a high-rate loading event, with the actual, measured hoop stretch magnitudes displayed by a distribution of radial rings. The maximum tensile hoop stretch occurs at the time when the bubble is largest, i.e. when *R*(*t*) = *R*_max_. We correspondingly write a quantity *λ_θ,_*_max_ as the hoop stretch experienced by cells at different radial positions at the time of maximum bubble expansion,

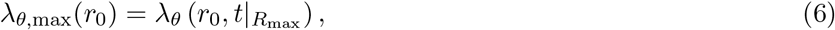

Thus, all hoop stretch levels *λ_θ,_*_max_ shown in Fig. 1 correspond to the maximum tensile stretches experienced by the cells as a function of radial distance from the bubble epicenter. Consistent with mechanics-based models of inhomogeneous deformations induced by an expanding and collapsing bubble, the deformation field decays rapidly with distance (~ 1/r^3^) away from the bubble wall.

As can be seen from the individual insets in Fig. 1d, there is a clear transition in cell morphology from a healthy network at stretches below *λ_θ,_*_max_ ≈ 1.05, to a pathology marked by significant morphological degeneration at stretches of *λ_θ,_*_max_ > 1.2 to a complete loss in Calcein AM signal at stretches of 2 and above. As fluorescence produced by Calcein AM indicates intact function of intracellular esterases of living cells, complete loss in Calcein AM signal indicates probable cell death. Spatial distances measured from the bubble center directly correlate with the amount of physical stretch experienced by the cells (see Eqs. 3, 4). Therefore, we use a simple image subtraction procedure of the Calcein AM signal in images before and after mechanical loading (i.e., cavitation). Notably, both of these images are taken of the reference configuration, but the difference between them allows us to quantitatively extract the radial boundary beyond which complete Calcein AM signal was present and below which it was lost. We denote this significant position in the tissue as *r*_0_ = *R*_PI_, which we call the primary injury radius.

Figure 2a illustrates the general analysis procedure for identifying critical injury thresholds associated with various pathological features shown in Figs. 1d and 3. Prior to cavitation-induced loading of each sample a 3D volumetric image is obtained as the healthy base, or reference, state. Next, we employ a modified form of our previously developed image fitting routines [24] to accurately extract the time-varying bubble radii, *R*(*t*), from each of the high-speed image frames recorded during the cavitation event. Finally, a high-resolution 3D post-scan of the same volumetric image stack is obtained and cellular morphologies (labeled via Calcein AM or various immunocytochemical markers) different from their respective base state are spatially marked.

**Figure 2:**
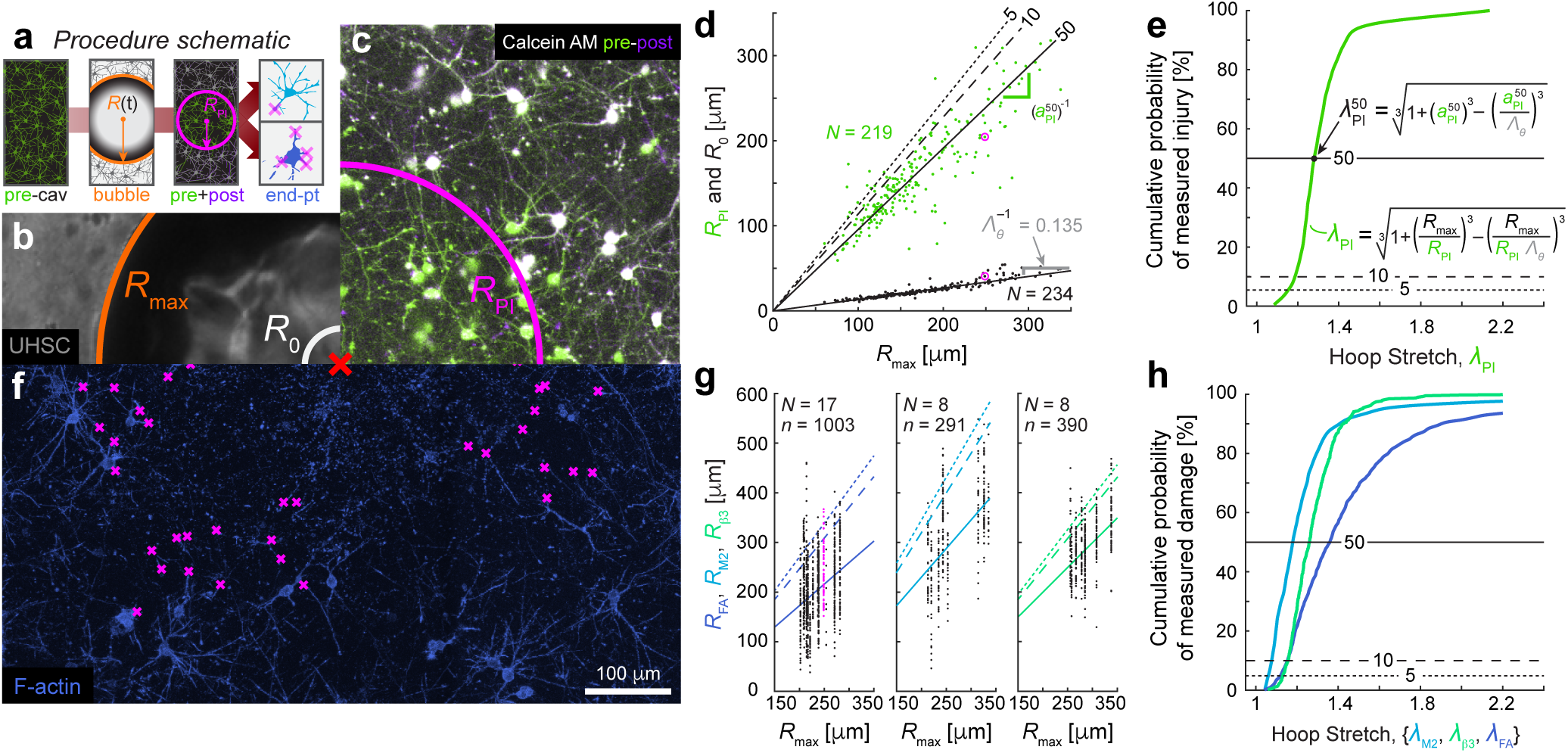
Procedure and image processing-based quantification of neural injury and critical stretch-based injury thresholds. (a) The mechanical history of a sample is defined by a chronological sequence of image sets: a 3D pre-cavitation reference set of live-stained cells, (b) a 2D ultra-high-speed video of the high-rate loading cavitation bubble dynamics, including maximum size *R*_max_ (orange) and quasi-equilibrium size *R*_0_ (white), and (c) a 3D post-cavitation image stack of live-stained neural cells. Maximum projections of the pre-(green) and post-cavitation (purple) image stacks are overlaid as an anaglyph image to illustrate the region of complete cellular damage (white color indicates the time-colocalization of cells before and after injury). A customized image subtraction algorithm computes a radius of primary injury (*R*_PI_, magenta) for each injured 3D volume. (d) Both the primary injury (PI, green, *N* = 219 experiments) and bubble equilibrium (black, *N* = 234) radii for each experiment (data points from (c) circled in magenta) scale linearly with maximum bubble radius, suggesting a constant and invariant critical strain value for primary mechanical injury. (e) Cumulative probability curve for defining the stretch-based thresholds for (or inverse tolerance of) primary injury based on Calcein AM (50%, solid; 10%, dashed; 5%, dotted). (f) For a subset of samples, end-point, 3D immunofluorescence imaging provided further insight into structure-specific injury thresholds. Degenerate and damaged projections were manually tagged within each sample (‘x’, magenta) through visual inspection. (g) Points were chosen for volumes (data from f highlighted in magenta) fluorescently labeled for f-actin (FA, royal blue; *n* = 1003 points over *N* = 17 volumes), MAP2 (M2, sky blue; *n* = 291, *N* = 8), and *β*3-tubulin (*β*3, spring green; *n* = 390, *N* = 8), converted into 3D radial measurements from the bubble center, and plotted versus the maximum bubble radius. (h) Cumulative probability curve for defining the stretch-based injury thresholds (or injury tolerance) for f-actin, *β*3-tubulin, and MAP2 (50%, solid; 10%, dashed; 5%, dotted). Scale bar for all images is 100 *μ*m.

Figure 2b provides a representative overlay of the best fit of the maximum bubble radius, *R*_max_, and the final bubble equilibrium radius, *R*_0_, onto the high-speed brightfield microscopy image. Across all performed experiments the values for *R*_max_ and *R*_0_ ranged from 60 − 338 *μ*m (*N* = 238) and 10 − 55 *μ*m (*N* = 219), respectively, with the ratio between them defining the maximum material stretch. As can been seen from Fig. 2d, the scaling between *R*_max_ and *R*_0_ is highly linear (bisquare *R*^2^ = 0.971) across a large distribution of maximum bubble radii with the median maximum bubble stretch ratio at the wall given by Λ_*θ*_ = *R*_max_*/R*_0_ = 7.40 (Fig. 2d, black).

For each experiment pre- and post-cavitation, Calcein AM volumes are compared using a custom image subtraction scheme in tandem with a quality-factor-based image correlation technique [28] to determine a quantitative radial assessment for primary injury, *R*_PI_ (Fig. 2c). Across all sampled values for *R*_max_, the primary injury radius-to-maximum bubble radius ratio, *a*_PI_ = *R*_PI_*/R*_max_ was direct (bisquare *R*^2^ = 0.869; *N* = 219) with median 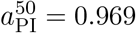 (Fig. 2d, green).

Due to the observation that injury occurs with less frequency further away from the bubble center, we construct an inverse injury tolerance curve, similar in spirit to the Wayne State Injury Tolerance Curve [29], for neural cells (Fig. 2e). Within the context of this study, injury is defined pathomorphologically using either Calcein AM or immunochemical markers for f-actin, *β*3−tubulin, and MAP2. Morphologically degenerate cellular features are identified in comparison to the healthy base state prior to mechanical loading and spatially marked with respect to the center of the cavitation bubble. Using Eqns. 1–4, we can define critical injury or damage stretch thresholds based on their position within the 3D image and relative to the stress-free, equilibrium bubble by,

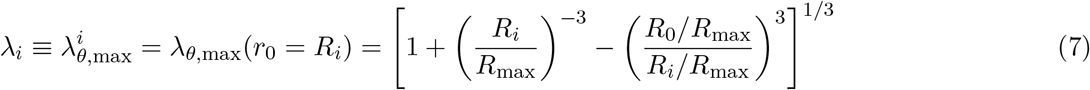

where *R_i_* corresponds to the radial position at equilibrium identifying morphologically degenerate cellular features within each 3D image (Fig. 2) for each of the employed structural markers, i.e., *i* = {primary injury (PI; Calcein AM), f-actin (FA), *β*3-tubulin (*β*3), MAP2 (M2)}. To reduce subsequent notation burden, we will henceforth write the maximum hoop stretch at a position 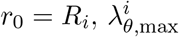, much more simply as *λ_i_*.

For example, to calculate the critical injury threshold for primary injury (characterized by a loss in Calcein AM signal, Fig. 2c) Eq. 7 can be recast in terms of Λ_0_ and *a*_PI_ using the information provided by Fig. 2d (i.e. the slopes of *R*_PI_*/R*_max_ and *R*_0_*/R*_max_) as,

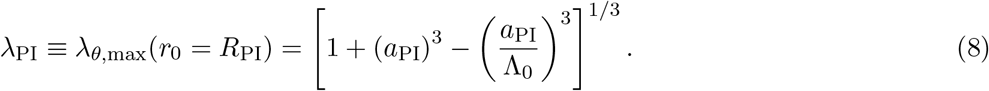

The specific value of a critical injury threshold can then, in general, be defined according to any desired percentile value within the cumulative distribution function; we report the stretch values corresponding to 5% (dotted), 10% (dashed), and 50% (solid) values as intuitive potential threshold options. These stretch values represent the stretches below which we only observe injury in the corresponding percentage of samples. For example, the 5% injury threshold signifies a stretch value below which only 5% of the samples sustained injury at lower stretches, or alternatively, 95% of samples were injured at stretches greater than the 5% threshold value. The [5-10-50]-th percentile thresholds correspond to stretch values, *λ*_PI_, can be found summarized in Table 1 and are on the order of ~ 1.2.

**Table 1:** Critical mechanical thresholds at [5,10,50]-th percentile thresholds. Critical stretch, hoop strain, hoop strain rate, and cellular stress estimates at the [5,10,50]-th percentiles of the cumulative injury function (Figs. 2 and 5) for each of the examined cellular and cytoskeletal damage features.

| Percentile: | Hoop Stretch |  |  | Hoop Strain |  |  | Hoop Strain Rate |  |  | Cellular Stress |  |  |
| --- | --- | --- | --- | --- | --- | --- | --- | --- | --- | --- | --- | --- |
| | $\lambda_{\theta} [-]$ | | | $E_{\theta\theta} [-]$ | | | $\dot{E}_{\theta\theta} \times 10^4 [\text{s}^{-1}]$ | | | $\sigma_{\theta\theta} [\text{kPa}]$ | | |
|  | 5<sup>th</sup> | 10<sup>th</sup> | 50<sup>th</sup> | 5<sup>th</sup> | 10<sup>th</sup> | 50<sup>th</sup> | 5<sup>th</sup> | 10<sup>th</sup> | 50<sup>th</sup> | 5<sup>th</sup> | 10<sup>th</sup> | 50<sup>th</sup> |
| *Primary Injury* | 1.152 | 1.187 | 1.287 | 0.141 | 0.171 | 0.253 | 1.143 | 1.424 | 2.259 | 2.404 | 2.871 | 4.101 |
| *f-actin* | 1.118 | 1.151 | 1.362 | 0.111 | 0.140 | 0.309 | 0.879 | 1.133 | 2.917 | 1.928 | 2.386 | 4.956 |
| *MAP2* | 1.061 | 1.075 | 1.182 | 0.059 | 0.073 | 0.168 | 0.448 | 0.557 | 1.384 | 1.062 | 1.294 | 2.807 |
| *$\beta 3$-Tubulin* | 1.132 | 1.153 | 1.261 | 0.124 | 0.143 | 0.232 | 0.987 | 1.154 | 2.029 | 2.128 | 2.421 | 3.781 |

To provide more detail on the pathological outcome of commonly reported structural constituents within the neural cells, we performed immunocytochemistry on a subset of samples. In contrast to the primary injury radii which are determined per sample (*N*), determination of critical injury thresholds for f-actin, *β*3−tubulin, and MAP2 are calculated at all manually tagged locations (*n*) within the immunofluorescence image of each sample (Fig. 2f). More details on the identification and tagging procedure is described in ‘Methods’. Analogous to calculating the primary injury critical stretch threshold, we calculate the critical point-wise stretch values via Eq. 7 for f-actin (*N* = 17*, n* = 1003), *β*3−tubulin (*N* = 8*, n* = 380), and MAP2 (*N* = 8*, n* = 291) using their respective radii ratios shown in Fig. 2g. As before, we construct a cumulative distribution function to determine critical structural injury thresholds for the [5-10-50]-th percentile of damaged projections for each marker (Figure 2h). The stretch values for the 5%, 10% and 50% injury thresholds of {*λ*_f-actin_, *λ*_MAP2_, *λ_β_*_3-tubulin_} are summarized in Table 1 and range from approximately 1 to 1.7.

To visually illustrate how well each of the above determined injury thresholds compares with representative micrographs for each of the identified structural markers, we overlay the calculated injury risk percentiles on maximum intensity projections of representative 3D volumes which experienced approximately equivalent maximum bubble expansion and hoop stretch (*R*_max_, Fig. 3). Figure 3a highlights the superposition of Calcein AM images taken before and after mechanical loading as an anaglyph image, while Fig. 3b–d are immunofluorescence images selecting for f-actin, MAP2, and *β*3-tubulin, respectively. Figure 3e–h highlights cells around the 5th, 10th and 50th percentile injury thresholds, with the primary injury radius *R*_PI_ highlighted by the magenta line, and several damaged projections highlighted by magenta triangles. Qualitatively, these images show that injury threshold percentiles between 10% and 50% provide a good lower (conservative) bound for predicting high-rate cellular injury.

**Figure 3:**
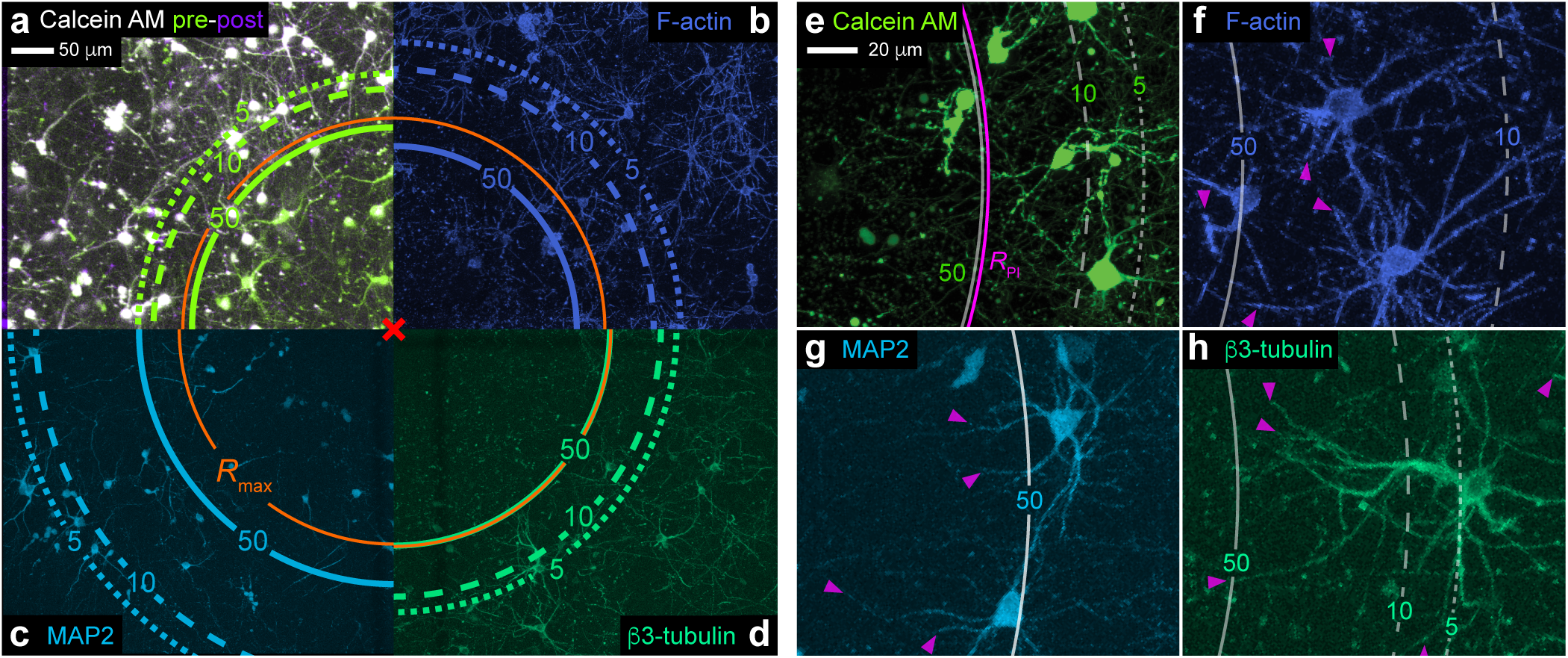
Representative maximum intensity projection micrographs of neural cell injury pathology and associated injury thresholds limits. (a) Anaglyph maximum projection image of pre-(green) and post-cavitation (purple) states of neural cells in 3D culture illustrating the damaging effect of a high-rate mechanical loading event; cells remaining healthy and intact are shown in white (indicating colocalization in both pre- and post-cavitation images). End-point maximum intensity projection images of immunocytochemical markers show (b) f-actin (royal blue), (c) MAP2 (sky blue), and (d) *β*3-tubulin (spring green) post-cavitation. For a-d, the bubble epicenter is located at the respective image corner (red ‘x’), and overlaid are the sample-specific *R*_max_ size (orange). Scale bar, 50 *μ*m. Expanded maximum projection micrographs of (e) live-stained neural cells (green) with (dataset-specific) associated primary injury radius (*R*_PI_, magenta) and cytoskeletal components (f) f-actin (royal blue), (g) MAP2 (sky blue), and (h) *β*3-tubulin (spring green) with (dataset-specific) tagged damaged projections (triangles, magenta) after high-rate mechanical loading. For (e-h), scale bar, 20 *μ*m. For all panels, overlaid are the corresponding 50% (solid line), 10% (dashed line), and 5% (dotted line) injury percentiles from the population analysis (Fig. 2), scaled by the sample-specific maximum bubble radii *R*_max_.

Up to this point, we have computed critical, stretch-based injury thresholds for Calcein AM, f-actin, MAP2, and *β*3-tubulin, and provided both a qualitative and quantitative assessment of the accuracy of these metrics in predicting structural (or primary) injury in our 3D *in vitro* model of TBI (Figs. 1–3). However, it is important to note that the deformations applied by the expanding and collapsing microcavitation bubble occur in a rapid, oscillatory fashion (Fig. 4a-b) subjecting cells to significant changes in the applied loading rate. Instead of expressing the rate of deformation in terms of a stretch rate, we utilize the widely used metric of strain rate.

**Figure 4:**
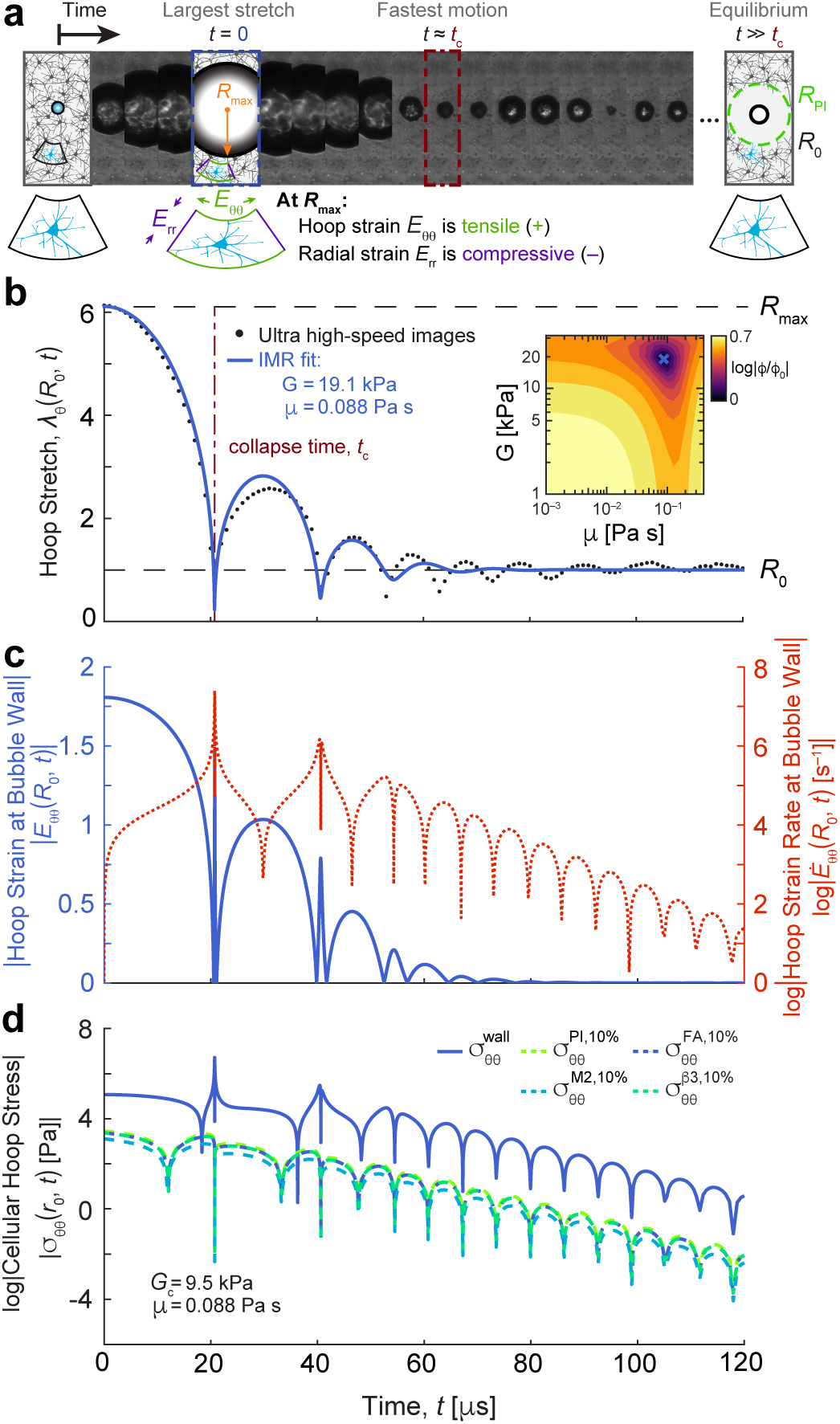
Temporal evolution of the applied and experienced cellular deformations and stresses. (a) High-speed time-lapse image series of a representative oscillating cavitation bubble generated inside the 3D neural tissue scaffold. *R*_max_ and *R*_0_ denote the maximum and equilibrium bubble radii, respectively, from which all relevant deformation quantities are calculated. (b) Generated material hoop stretch at the bubble wall, *λ_θ_*(*R*_0_*, t*), due to the microcavitation bubble. Black dots indicate experimental data points extracted from high-speed videography to compute hoop stretch. Solid blue lines denote best fit, demonstrated by a defined minimum in the least squares error cost function space *ϕ*(*G, μ*) (inset), of the cavitation dynamics using our previously developed IMR technique [24] and assuming the neural tissue to be well-described by a non-linear Kelvin-Voigt material model (See Methods). (c) Temporal evolution of the absolute magnitude of the logarithmic (Hencky) hoop strain, *E_θθ_*(*R*_0_*, t*) (blue line), and the temporal evolution of the log of the magnitude of the hoop strain rate, 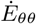 (dashed orange line), each computed from the IMR best-fit material properties in (b). (d) Absolute cellular hoop stress estimates over time, assuming a nonlinear Kelvin-Voigt constitutive model with *G* = 9.5 kPa and *μ* = 0.088 Pa*·*s, at the bubble wall (solid blue) and spatial locations corresponding to the 10% damage thresholds for primary injury (PI, dashed green), *β*3-tubulin (*β*3, dashed spring green), MAP2 (M2, dashed sky blue), and f-actin (FA, dashed royal blue).

The advantage here is that both strain rate and strain are commonly-employed quantification metrics of mechanical deformations used in cell mechanics, biophysics [6, 30–33] and computational modeling of brain injury [22, 34–36]. Conveniently, the logarithmic, or Hencky, strain is simply defined as the natural log of the material stretch, i.e., *E_θθ_* = *E_ϕϕ_* = ln(*λ_θ_*), and *E_rr_* = −2 ln(*λ_θ_*), where both strain and stretch are functions of position *r*_0_ and time *t*. As neural cells are commonly considered to be susceptible to tensile deformation [34, 37–40], we concern ourselves primarily with the tensile hoop strain components *E_θθ_* = *E_ϕϕ_*, just as we did in the case for the hoop stretch, *λ_θ_*. The strain rate is then the material (or total) time derivative of the material hoop strain, 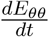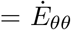.

Figures 4b and 4c provide a general overview of the dynamic changes in hoop stretch (Fig. 4b), hoop strain and strain rate (Fig. 4c) at the bubble wall, which is the location of maximum stretch and strain generated in the material. One apparent feature in each of the plots in Figures 4b and 4c are the sharp curvature points around the various bubble collapse points (e.g., *t* ≃ 20*μs,* 40*μs*, etc.), which produces some of the highest loading rates observed in the material 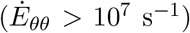. To fully resolve these transitions in bubble and material behavior we utilize our previously developed inertial microcavitation rheometry (IMR) technique, which allows for both least squares fitting and temporal interpolation of the entire cavitation process.

During maximum expansion, logarithmic hoop strains near the bubble wall are large and finite, peaking at *E_θθ_* = 1.81 for the representative sample in Figure 4c (blue curve). Logarithmic strain rates (Fig. 4c, red dotted curve) are similarly large, reaching peak values of 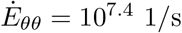 at the time of bubble collapse, *t_c_*. At all times, the applied strain rates as shown in Fig. 4c are significantly higher than those commonly reported in the blunt and mild TBI literature [6, 31, 41, 42], consistent with high-loading rate phenomena encountered in blast, cavitation or pressure-driven (sonic) scenarios. Analogous to our stretch calculations, we compute the time-maxima of the hoop strain due to the cavitation bubble, 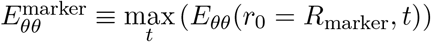, and of its associated strain rate, 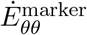 at the percentile damage threshold locations. The critical hoop strains, 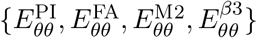, and maximum strain rates, 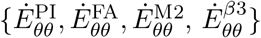 at the [5-10-50]-th percentile damage threshold locations are summarized in Table 1 and are on the order of 0.06 to 0.25, and ~ 10^4^s^−1^, respectively.

Since neural cells, like many other bodily tissues, are well-described as rate-dependent viscoelastic materials, one way to account for both the large strains and high strain rates experienced by the cells is to introduce a rate-dependent, large deformation constitutive model for estimating actual cell stresses. Such an approach is quite common for biological scenarios that show strong loading rate dependence [43, 44]. To estimate the deviatoric (i.e. removing the effect of the atmospheric baseline pressure, *p_∞_*) cellular stresses we employ a simple, non-linear Neo-Hookean Kelvin-Voigt model capable of both describing the large strain elastic, and high-rate viscous behavior. Furthermore, based on our prior work [6], we assume displacement continuity across the cell-collagen gel interface, allowing us to prescribe the same strains and strain rates to the cells as measured by IMR in our neural tissue model (Fig. 4c). The deviatoric hoop stress for a non-linear Kelvin-Voigt material takes the form of:

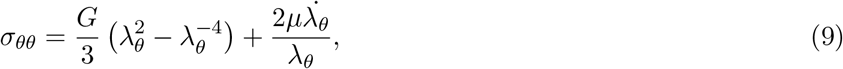

where *G* and *μ* denote the dynamic shear modulus and viscosity of the material, respectively. It can be seen from Eq. 9 that the total cell stress in the Kelvin-Voigt model is a straightforward addition of the elastic and viscous stress terms.

To compute cellular stresses we estimate both *G* and *μ* from previously published literature estimates. The elastic properties for neural cells can vary based on the examined subcellular compartment, with somatic values approaching several hundred Pa [45] and stiffness estimates of axonal projections ranging from 9.5-12 kPa [46, 47]. Considering that most of our pathology overwhelmingly presents degeneration and damage along cytoskeletal projections, we adopt *G* = 9.5 kPa as our elastic parameter for estimating the mechanical stresses within neural cell projections. Given a scarcity of viscosity measurements for cells, particularly for neural cells under high loading rates, we estimate the dynamic viscosity, *μ*, for our neural cells using a combination of material calibration measurements via IMR, and a scaling argument based on literature values. First, using IMR we determined the dynamic viscosity of our 3D collagen-cell *in vitro* model to be approximately 0.088 Pa·s (Fig. 4b). Since the volume fraction of neural cells within our collagen scaffold is less than one percent, we assume that the viscous behavior is predominantly driven by the collagen fibers. When comparing our high-rate, dynamic viscosity measurement to low, or quasi-static rate (~ 0.1–1 Hz) estimates from the literature for type I collagen gels, i.e., values around ~ 2–3 Pa·s [48, 49], we, consistent with the literature, attribute this discrepancy to the well-known shear-thinning effect of collagen at high-rate [50, 51]. Since quasi-static, low frequency estimates on the dynamic viscosity in neurons have been reported to be around ~ 2–3 Pa·s [52], which is very similar to type I collagen at low frequencies, we assume a similar scaling on the high-rate dynamic viscosity as we did with collagen, and thus assign the same value of our cell-gel system to the cells directly, i.e., *μ* = 0.088 Pa·s.

Figure 4d displays the temporally evolving cytoskeletal cell stresses at the bubble wall (solid blue), as well as at the locations corresponding to the 10th percentile injury probability for each of the cytoskeletal structures (dashed). Estimates of the maximum stress experienced by cells at positions of interest, 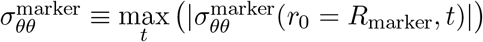, corresponding to the [5,10,50]-th damage thresholds for 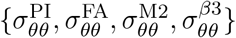, are summarized in Table 1 and range from ~1-5 kPa. Since the cellular stresses are a sum of the rate-dependent viscous and rate-independent elastic stress contributions, the shape and evolution of the cell stresses present a temporal shape evolution in between *E_θθ_* and 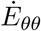 (Fig. 4c), with the highest stresses occurring closest to the bubble wall (~ 5.2 MPa).

## Discussion

Our current ability to predict, diagnose, and mitigate high-rate tissue injuries such as those that arise during advanced, sonic- and laser-based medical treatments, or in hazardous environments including blast exposures, is limited by our understanding of the unique pathology of high-rate injury itself and the resolution of associated critical injury thresholds. By carefully integrating a physiologically relevant, 3D *in vitro* neural tissue culture model with our recently pioneered inertial microcavitation rheology (IMR) technique, we provide high-resolution, structural information on the unique injury pathology of cells experiencing high-rate mechanical loading, i.e., strain rates in excess of 10^3^ 1/s. Using 3D volumetric image reconstruction via confocal and multiphoton imaging in tandem with IMR, we provide quantitative estimates of several key physical quantities producing this pathology (Fig. 5). Specifically, we provide estimates of critical stretch, strain, strain rate and cell stress thresholds for the failure of neural projections containing common subcellular structural components such as, microtubules, filamentous actin (f-actin), and microtubule-associated protein-2 (MAP2).

**Figure 5:**
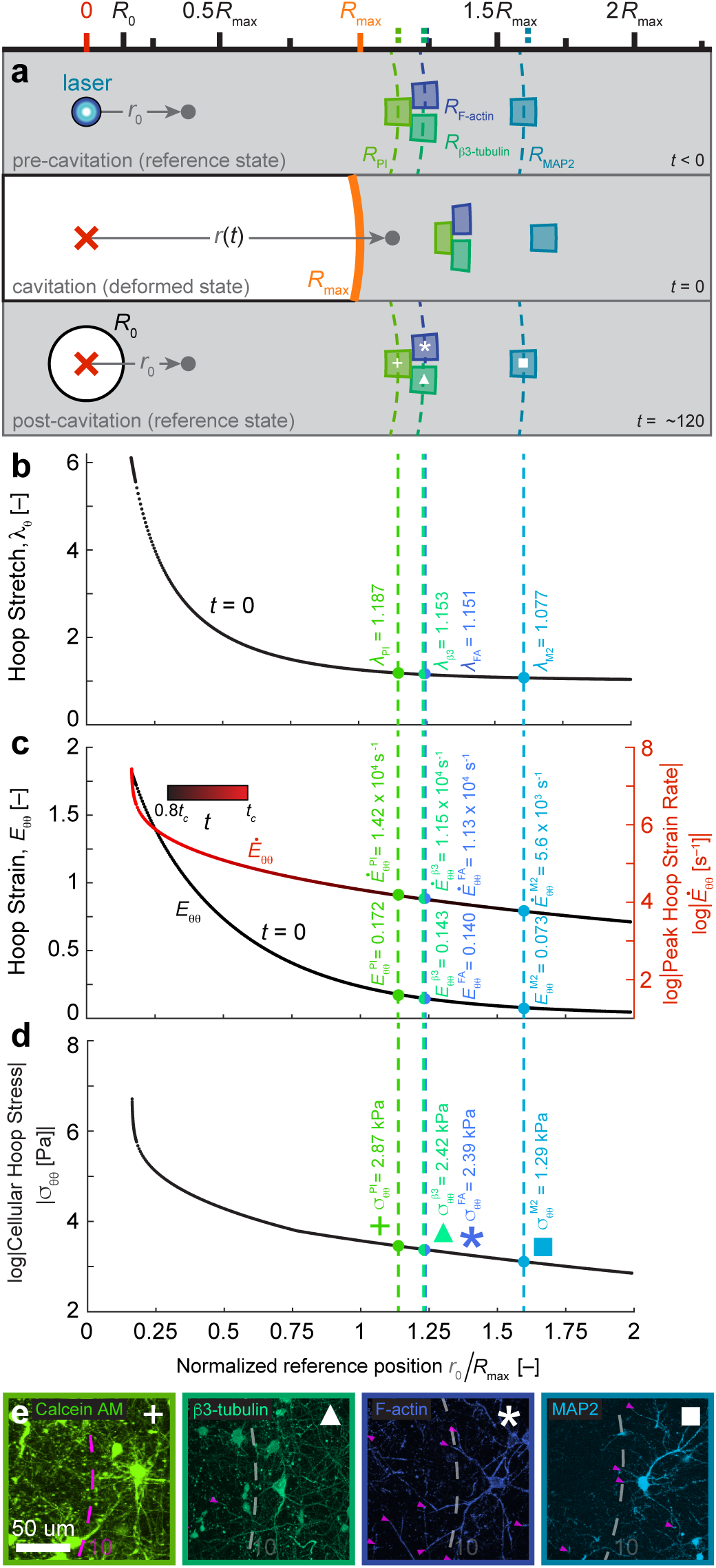
Critical injury thresholds for neural cells in terms of stretch, logarithmic (Hencky) strain and strain rate, and cellular stress as a function of normalized position *R*_0_/*R*_max_. (a) Shown is a to-scale three-panel schematic of our neural tissue just before cavitation (*t* < 0), at maximum bubble size (*t* = 0), and at quasi-equilibrium post-cavitation (*t* ≫ 0). Highlighted are representative volumes at the true spatial locations of the 10th injury percentile for primary injury (PI; green plus), and degenerate projections for *β*3-tubulin (*β*3, spring green, triangle), f-actin (FA royal blue, star), and MAP2 (M2, sky blue, square). During maximum bubble expansion volumes are displaced and stretched, with volumes closer to the bubble wall experiencing higher strain than those farther away. (b) Hoop stretch, *λ_θ_*, as a function of normalized radial distance within the neural tissue at time of maximum bubble expansion/tissue stretch and to-scale with (a). Spatial locations corresponding to the 10th percentile injury stretch for primary injury (green) and cytoskeletal components *β*3-tubulin (spring green), f-actin (royal blue), and MAP2 (sky blue) are *λ_θ_* = {1.187, 1.155, 1.154, 1.077}, respectively. (c) Hoop strain at maximum bubble expansion (black) and strain-rate (red to black gradient) as a function of position in the material during the full dynamic cavitation process. Values of hoop strain corresponding to locations of the 10th injury percentile for primary injury (green) and cytoskeletal components *β*3-tubulin (spring green), f-actin (royal blue), and MAP2 (sky blue) are *E_θθ_* = {0.172, 0.143, 0.141, 0.073}, respectively. The maximum strain rates experienced at the same locations are 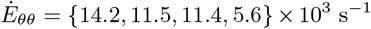, respectively, and occur at times *t* at, or slightly before, the time of bubble collapse, *tc*. (d) Cellular hoop stresses as a function of position corresponding to the positions of 10th percentile injury had corresponding values *σ_θθ_* = {2.87, 2.39, 1.29, 2.42} kPa, respectively, during the full dynamic cavitation process. (e) Representative fluorescent maximum intensity projections of injured cells at positions of the 10% damage thresholds for primary injury (plus), *β*3-tubulin (triangle), f-actin (star), and MAP2 (square).

In particular, we observe differing stretch tolerances of projections containing different subcellular components, which can be found summarized in Table 1 and Fig. 5. In analyzing the relatively conservative 10th percentile stretch damage thresholds we find that, in order of decreasing stretch sensitivity, projections containing MAP2 are damaged at the lowest strains of 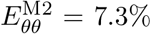, followed by f-actin- and *β*3-tubulin-containing projections at strains of 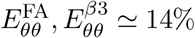 (Figure 5c). The total disruption of overall cellular architecture (*R*_PI_)—specifically the depletion of intracellular esterase activity—was found to require the largest strains of 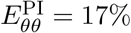.

The selection of the cytoskeletal markers examined here are intended to provide granularity with respect to the susceptibility of cell-type and neuronal sub-compartments to high-rate, stretch-induced damage. Given the nature of our polyculture of neuronal and glial cells used here, the selection of *β*3-tubulin for microtubules provides specificity for neuronal projections, whereas f-actin is general to the full neural cell population (including astrocytes) providing a more general projection failure criteria across all cell-types [7]. The measured failure of f-actin- and microtubule-containing projections, in aggregate, under nearly equivalent loading, is intuitive at diffraction-limited length scales for the following reason. F-actin and microtubules are co-localized throughout the majority of the cytostructure of most neural cells despite having different intracellular organization and persistence lengths based on the sub-compartment and cell type examined [38, 39, 53]. For example, in the axon of a neuron, f-actin forms periodic ring structures with spectrin and other proteins, while microtubules form long aligned structures along the length of the projection [53]. Both f-actin and microtubule components have been shown to exhibit a rate-dependent susceptibility *in silico* to tensile failure in neural cells with an increased dependence on the rate of deformation [38–40]. These high-rate (10^8^ − 10^9^ 1/s) molecular dynamics (MD) simulations indicate that onset of failure for microtubules and actin-spectrin structures begins at different logarithmic strain thresholds of ~ 0.15 and ~ 0.17, respectively [38, 39]. The closeness of our measured 10th percentile stretch damage thresholds (at strains of ~ 0.14) for both f-actin and microtubules, coupled with the MD prediction of different failure criteria for f-actin and microtubules, suggests that in the combined system microtubules may act as the limiting component of the projection structure. The damage of projections containing MAP2, which is generally consigned to microtubules of the dendrites of mature neurons [54], at lower strain thresholds (~ 0.07) than general neuronal microtubule-containing (~ 0.14 strain) projections suggests the existence of a dependence on the micro-structural differences of axons vs. dendrites [53] with respect to strain tolerance. We thus interpret the apparent failure of MAP2-containing, or dendritic, projections (under equivalent loading) at a lower strain threshold as the potential existence of a stress-dependent injury criterion.

The existence of a stress-dependent injury criterion can be intuitively understood through the notion that a load exerted on a structure will be experienced differently depending on the organization and interaction between the structural components within the system. A more nuanced interpretation comes from an understanding of the role the organization and structure of the cytoskeleton of different subcellular compartments play in the cells’ susceptibility to injury. The arrangement of microtubules differs between neuronal compartments. Axons can consist of ~ 10 − 100 microtubules organized into uniform bundles with the plus-end oriented toward the synapse; these bundles are stabilized through binding with tau proteins [55], whereas in dendrites the microtubule orientation is diverse, with many microtubules oriented with the minus-end oriented away from the cell body stabilized by MAP2 [54, 55]. Similarly, the organization of f-actin in the axon and dendrites differ. In the axon, f-actin and spectrin form periodic rings along the length of the axon, whereas the dendrites consist of a mix of linear and branched f-actin networks [53, 56]. The rate dependence for failure of these structures has been shown to relate to protein associations within the structure, including the susceptibility of spectrin unfolding under high-rate loading for f-actin failure [39], and sliding-stretching behavior of microtubules through the viscous unfolding and dissociation of microtubule-associated proteins [57]. The rate-dependent protein association and micro-scale structural protein architecture differences signal the importance of mechanical thresholds that take into account the complexity of the stresses experienced at locations of cellular failure. To address this, we provide initial estimates of the cellular stress for neural projection failure (Fig. 5c), as summarized in Table 1 at the [5-10-50]-th stretch damage thresholds. While these estimates incorporate the non-linear elastic, and linear viscous/rate-dependent contributions of the physical structure of each neural cell, they only provide whole system (i.e., the entire cellular projection modeled as one) stress estimates, rather than stress estimates on the actual subcellular structural units. Resolution of subcellular, protein-level-specific injury stresses is an exciting area of active research, and current modeling efforts should shed more light onto the distribution of internal stresses within the overall structure-level stresses presented here. The representative morphology of cellular damage metrics is highlighted at the 10th percentile damage thresholds in Figure 5e, where example degenerate projections can be seen highlighted.

Lastly, we note that all of the observed cellular injury falls within 2.5 times of the maximum bubble radius for each data set (Fig. 1d), beyond which all strains and elastic stress contributions become negligible (e.g., *E_θθ_* < 2.5%) Fig. 5c-d. Interestingly, the strain rate remains non-zero (Fig. 5c) leading to a small remnant deviatoric viscous stress (< 400 Pa), which we do not observe to be injurious. These spatial scaling observations are consistent with our previous analysis of inertial cavitation bubbles showing that most of the significant material deformations occur within 2.5 times *R*_max_ from the bubble center [24].

While our work has focused on the acute pathology and structural metrics of injury, future work will investigate potential additional injury sequelae often seen in cellular and tissue injuries, including, but not limited to, the inflammatory response and activation of glial cells including astrocytes and microglia. One of the significant advantages of the high-rate loading model presented here is its optical nature and modularity, that is, it can be straightforwardly adapted to introduce high-rate loading in tissues of heightened physiological relevance including cortical spheroids, organoids, and *in vivo* tissues [8,58]. It should be noted that while our cavitation-based approach is minimally invasive and modular, the nature of the applied deformations are intrinsically tied to the resulting cavitation bubble dynamics, often introducing an oscillatory, repeat loading pattern of spatiotemporally varying strain and strain-rate. This is a deviation from much of the previous literature on TBI, in which most experimental platforms subjected cells and tissues to relatively simple, single, and uniform deformation profiles [6, 25, 30, 31, 41]. However, one could argue that the cyclic loading profile of the oscillating bubble reflects a more physically realistic loading scenario during real-world events or in clinical applications. [8].

Our IMR technique is not just limited to investigations detailing neural injury, but rather can be employed broadly across many different tissue models and anatomical constituents. Finally, while IMR provides a highly controlled and repeatable means of introducing high-rate mechanical loading to cellular and tissue constructs, the theoretical backbone of IMR also allows for the extraction of relevant high-rate material properties from the tissue system under interrogation. In the case for our collagen-based neural cell tissues we showed that using a non-linear Kelvin-Voigt material model, IMR determined the dynamic shear modulus and viscosity to be *G* = 19.1 kPa, and *μ* = 0.088 Pa·s, which falls within the the range of prior characterization results on brain and brain-like tissues [59, 60]. Thus, the IMR framework allows for the inverse determination of the physical properties (for example, the elastic shear modulus, dynamic viscosity, and analogous parameters for other viscoelastic models [65]) for various tissues and cellular systems at high-rate (e.g., ballistic and blast loading rates), which remain to be a significant challenge to obtain via traditional characterization methods. Coupled with data-driven methods for constitutive model identification [61], IMR has considerable potential for high-rate material calibration for these complex biomaterial systems.

In summary, in this study we provide significant new insight into the cellular pathology of neural cells undergoing high-rate deformations such as those arising in advanced sonic and laser-based medical procedures or hazardous environments that feature significant internal or external pressure exposures (e.g., directed-energy or blast exposures). Specifically, we provide quantitative information on the critical stretch, strain and stress values required to injure neural cells during high rating exposures, which should provide a scientific basis for the development of advanced brain injury predictive diagnostics, and mitigation strategies in medical procedures to limit collateral tissue damage, especially in the central nervous system.

## Methods

### Isolation of primary neural cells

All procedures were performed in full accordance with the Institutional Animal Care and Use Committee’s of Brown University and the University of Wisconsin, Madison. Unless otherwise stated, reagents were acquired from Thermo Fisher (Waltham, MA). Primary neural cells were isolated from postnatal day 0-1 Sprague-Dawley rats (Charles River, Wilmington, MA). Pups were anesthetized with hypothermic treatment and euthanized through cervical dislocation. The cerebral hemispheres were removed and placed in a cold solution of Hibernate A (HA; Brain Bits LLC, Springfield, IL) + 2% B-27 serum-free supplement (50×) + 0.25% GlutaMAX (100×), ventral side down. A cut was made around the lateral side of the brain, just above the entorhinal and perirhinal cortices, followed by separation at the corpus callosum. The excess tissue was removed from the cerebral cortex and the cortices were minced (~0.5 mm) and digested in a solution of Hibernate A (HA)–CaCl_2_ (Brain Bits LLC) + Papain (2 mg/mL; Worthington, Biochemical Corporation, Lakewood, NJ) for 30 minutes at 37°C with gentle agitation every 5 minutes. An equal volume of pre-warmed HA/B-27 was added to the HA-CaCl_2_/papain solution and centrifuged at 150g for 5 minutes. After removal of the supernatant, the dissociated tissue was re-suspended by gentle mechanical trituration in cortical complete media (Neurobasal-A medium + 1% Penicillin/Streptomycin + 0.25% GlutaMAX 100×), allowed to settle for 1-2 minutes, and passed through a sterile cell strainer (40 *μ*m mesh size) to filter large tissue debris. The suspension was again centrifuged at 150g for 5 minutes, the supernatant was removed, replaced with 3 mL cortical complete media, and cells were re-suspended by gentle mechanical trituration and counted with a hemocytometer (Hausser Scientific, Horsham, PA). Cell solution was diluted with cortical complete media to 10^6^ cells/mL prior to preparation of 3D collagen gels to yield a final seeding density of 3,750 cells/mm^3^ [6, 62, 63]. Additional information and specifics of the cortical isolation and a more detailed protocol can be found in Scimone et al [7].

### Preparation of 3D collagen hydrogels

Type I Collagen (rat tail; Dow Corning, Tewksbury, MA) hydrogels were made at a concentration of 2.2 mg/mL according to the manufacturer’s standard protocol. 10× Phosphate-buffered Saline (PBS) was mixed with type I collagen stock solution (3.3−3.7 mg/mL) in 0.02 M acetic acid, triturated, and kept on ice. Neuron-cell media suspension was then added to the collagen mixture to get a collagen concentration of 2.2 mg/mL and neuron density of 3,750 cells/mm^3^, and brought to a target pH of 8.4 through the addition of 1 M NaOH (Sigma-Aldrich, St. Louis, MO). Collagen gel composition is shown in Table 2.

**Table 2:**
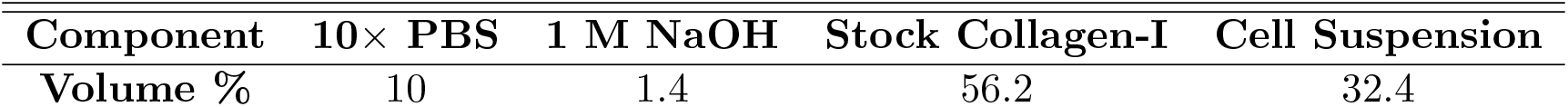
Concentrations for 3D type I Collagen hydrogels.

| **Component** | **10× PBS** | **1 M NaOH** | **Stock Collagen-I** | **Cell Suspension** |
| --- | --- | --- | --- | --- |
| **Volume %** | 10 | 1.4 | 56.2 | 32.4 |

**Table 3:** Immunolabeling probes for proteins of interest

| Antibody Target | Species | Manufacturer | Catalog No. | Concentration |
| --- | --- | --- | --- | --- |
| *Primary Antibodies* |  |  |  |  |
| β3-tubulin | Mouse (IgG) | ThermoFisher Sci. | MA1-19187 | 1:200 |
| MAP2 | Rabbit (IgG) | ThermoFisher Sci. | PA5-17646 | 1:200 |
| *Secondary Antibodies* |  |  |  |  |
| Anti-Mouse AF555 | Goat | ThermoFisher Sci. | A32727 | 1:100 |
| Anti-Rabbit AF555 | Goat | ThermoFisher Sci. | A32732 | 1:100 |
| *Phalloidin Probes* |  |  |  |  |
| Filamentous actin |  | ThermoFisher Sci. | R37112 | 4 drops/mL |

50 *μ*L volumes of cell-collagen gel solution were seeded directly into 48-well #1.5 glass-bottomed plates (MatTek, Ashland, MA). Dimensions of the samples were approximately 6 mm in diameter and 1.2 mm in height. Gels were polymerized in a humidified incubator at 37°C in 5% CO_2_ for approximately 20 minutes after which 300 *μ*L of cortical complete media is added to each well after polymerization. Each plate of samples was incubated at 37°C, 5% CO_2_ for 7 days *in vitro*, after which neurons showed a high degree of network connectivity (Fig. 1a).

### Live-cell fluorescence

Cell morphology was assessed before and immediately following microcavitation via cell-permeant Calcein Acetoxymethyl (AM). Cell cultures were incubated with 200 *μ*L of pre-warmed HA+2% B-27 buffer with Calcein AM (20 *μ*M) for 45 minutes to 1 h to allow for sufficient penetration into the gel, after which media was replaced with naïve pre-warmed HA+B-27 buffer. Samples not exhibiting healthy, fully-connected morphology were discarded from further data processing. Control cultures were incubated with 200 *μ*L of pre-warmed HA+2% B-27 buffer containing both Calcein AM (20 *μ*M)for 1 hr, after which media was replaced with naïve pre-warmed HA+B-27 buffer.

### Fixation and Cytoskeletal Labeling

Neural cell cultures were fixed post-live-cell fluorescent imaging using a solution of 4% v/v paraformaldehyde (Electron Microscopy Sciences, Hatfield, PA) + 8% w/v sucrose in 1× PBS for 30 minutes. Samples were subsequently washed four times with 1× PBS and stored at 4°C prior to additional labelling procedures. Samples were permeabilized in 0.5% (v/v) Triton ×100 for 30 minutes, washed three times with 1× Phosphate Buffered Saline (PBS). Non-specific staining was limited through a blocking step using 10% (w/v) Bovine Serum Albumin (BSA) (Jackson ImmunoResearch, PA, USA) for 2 hours at room temperature (measured at 22-23°), and all subsequent wash and stain steps were completed using 0.5% (w/v) BSA at room temperature. After blocking, samples were washed twice and stained overnight at room temperature for the primary antibody marker of interest, see Table 2. Samples were washed four times, and submerged in secondary antibody for six hours. Alternatively, if not antibody-stained, samples were washed four times and stained with Phalloidin stain ActinRed 555 ReadyProbes for 2 hours at 4 drops/mL in 0.5% BSA and washed four times with 1× PBS.

### Microscopy and imaging

For all live-cell experiments and controls, three-dimensional image stacks of cells immediately before and immediately following laser-induced cavitation were acquired using a Nikon A-1 laser scanning confocal or multiphoton system, mounted on a Ti-Eclipse inverted optical microscope controlled by NIS-elements software (Nikon, Tokyo, Japan). To maintain physiological conditions, a custom-made environmental chamber was built around the microscope and thermally-controlled at 37°C with a closed-loop Air-Therm heater (World Precision Instruments, Sarasota, FL). Samples in 48-well glass-bottomed plates were secured using a well plate holder (Ti-SH-W; Nikon), which was allowed to thermally equilibrate before imaging to avoid thermal drift in imaging locations. All cavitation events were performed 600 *μ*m above the top of the glass coverslip, or approximately at the mid-plane of the gels.

### Laser-induced microcavitation

Experimental inertial cavitation was generated via single pulses of a 3-5 ns frequency-doubled 532 nm Q-switched Nd:YAG laser (Continuum, San Jose, CA) through a Plan Fluor 20×/0.5 NA objective (Nikon Instruments, Japan). The beam was redirected into the back aperture of the objective by a 532 nm notch dichroic reflector (Semrock, Rochester, NY), and expanded to fill the back aperture of the imaging objective using a variable beam expander (Thorlabs, Newton, NJ). Upon passing through the objective, pulses converged at the image plane as validated by a continuous exposure alignment of a 635 nm laser diode (LDM635; Thorlabs, Newton, NJ) and were imaged using bright-field microscopy at either 270,000 fps on a Phantom V2511 (Vision Research, Inc., NJ, USA) or at 1,000,000 fps on a Shimadzu HPV-X2 (Shimadzu Corporation, Kyoto, Japan). The condenser aperture was reduced as much as possible without overly reducing image intensity to minimize effects of non-parallel light interacting with the bubble. As each 3D collagen gel had dimensions of 3 mm radius and 1.2 mm height, and to reduce potential biological effects of cellular death factors and mechanical effects of introduced boundaries, only one cavitation event was generated in each non-control sample at the approximate middle z-plane of 600 *μ*m.

### Cavitation Bubble Determination

**Algorithm 1:**
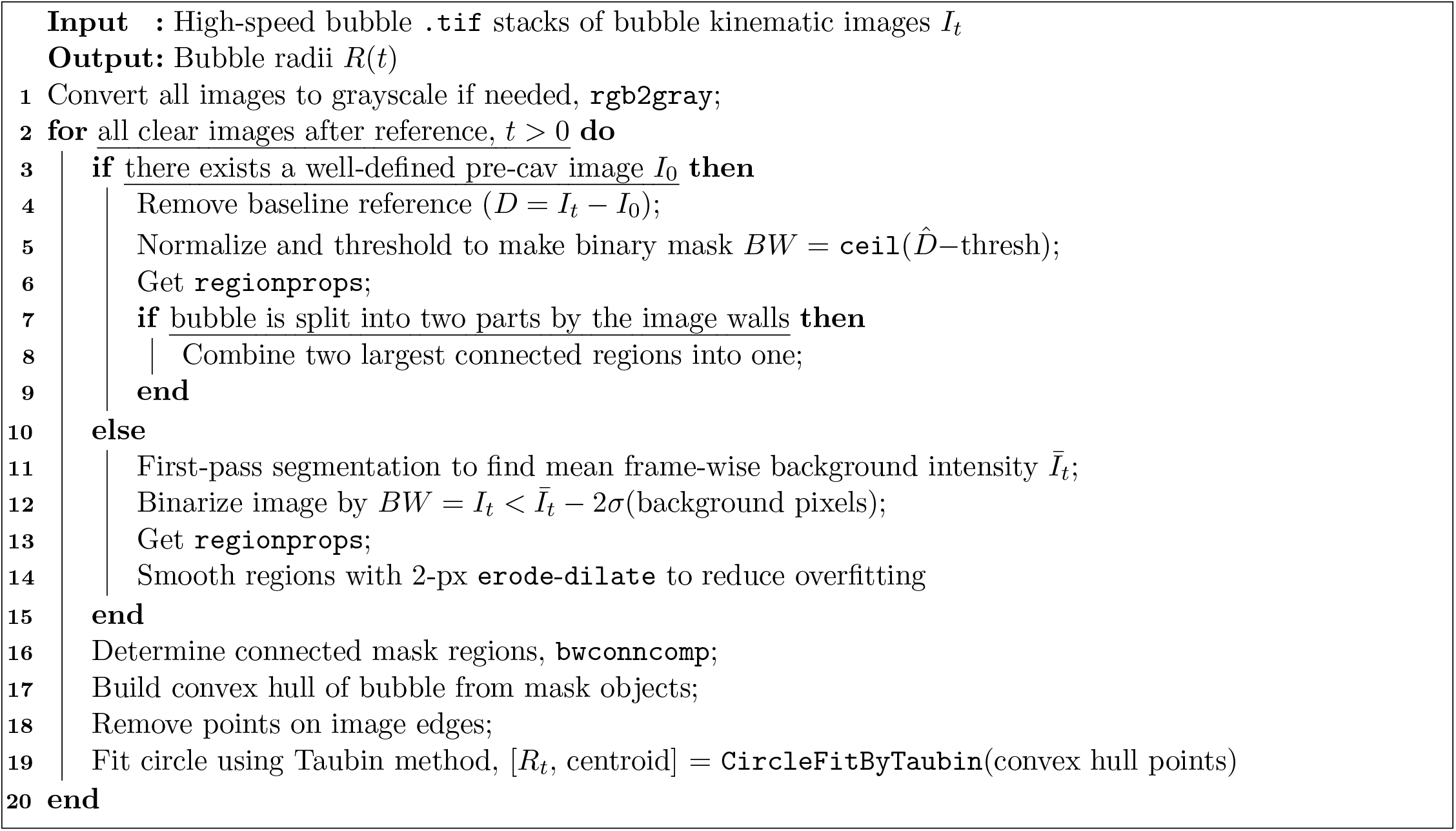
Calculating *R_max_* and *R*_0_ from high-speed bubble images.

The general processing procedure for acquiring the maximum bubble radius and equilibrium radius from high speed video is outlined in Algorithm 1. Each time-series of images consisted of either 101 frames of 512×128 px from a Phantom v2511 high speed camera, or 256 frames of 400×250 from a Shimadzu HPV-X2 high speed camera. Imaging frame rates were set to 270k fps (3.7 *μ*s/frame) for the Phantom v2511 and 1M fps (1 *μ*s/frame) for the Shimadzu HPV-X2 with a calibration image of an etched calibration grid taken for each to set the *μ*m/pixel ratio for each time-series (v2511, 1.38*μ*m/pixel; HPV-X2, 1.62*μ*m/pixel). For cases where there was a well-defined reference image, the first image in each time-series was designed as a reference image *I*_0_ of the gel before arrival of the laser pulse. For subsequent images of the bubble *I_t_*, differential images *D* = *I_t_* − *I*_0_ were constructed, normalized, and thresholded to create binary masks. In cases without a reference image, a first-pass segmentation defined a background intensity threshold to create binary masks. The built-in function bwconncomp (Image Processing Toolbox, Matlab) was then used to segment the bubble area. A convex hull was constructed around the bubble object excluding points on the image boundaries. Finally, points along the convex hull were then fit using a generalized eigenvector fitting method developed by Taubin [64] for the specific case of a circle. Bubble images were overlaid with the circle fit outline and manually inspected to ensure accurate fitting.

### 3D Image acquisition

Prior to laser-induced microcavitation, reference image stacks of 1024×1024×11 px (1.24×1.24 *μ*m spacing in x-y, 10 *μ*m spacing in z) images of Calcein AM-stained neural cells were acquired with a 10×/0.3 NA Plan Fluor objective (Nikon) centered at 600 *μ*m above the top surface of the bottom coverslip. Rather than create a full 3D reconstruction, total volume size and laser settings were chosen to ensure minimal effects from phototoxicity, resulting in a pseudo-3D confocal volume stack. Following initial imaging, cavitation was generated as described above, consistent with prior work [24, 65]. Samples were imaged and cavitated as described above in sets of 6-8 to reduce the time delay between cavitation and post-imaging. After completing a set of samples, the dichroic was removed and the 10×/0.3 NA Plan Fluor objective was repositioned to acquire a post-cavitation stack of images, again with dimensions 1024×1024×11 px (1.24×1.24 *μ*m in x-y, 10 *μ*m spacing in z). Laser scanning multiphoton imaging of immunolabelled subcellular structures was performed using a 25×/1.1 NA Apochromat water immersion objective (Nikon Instruments, Japan). Multiphoton image stacks of 1024×1024×113 px (0.5×0.5×1.35 *μ*m spacing) were taken post-fixation for structural regions of interest.

**Algorithm 2:**
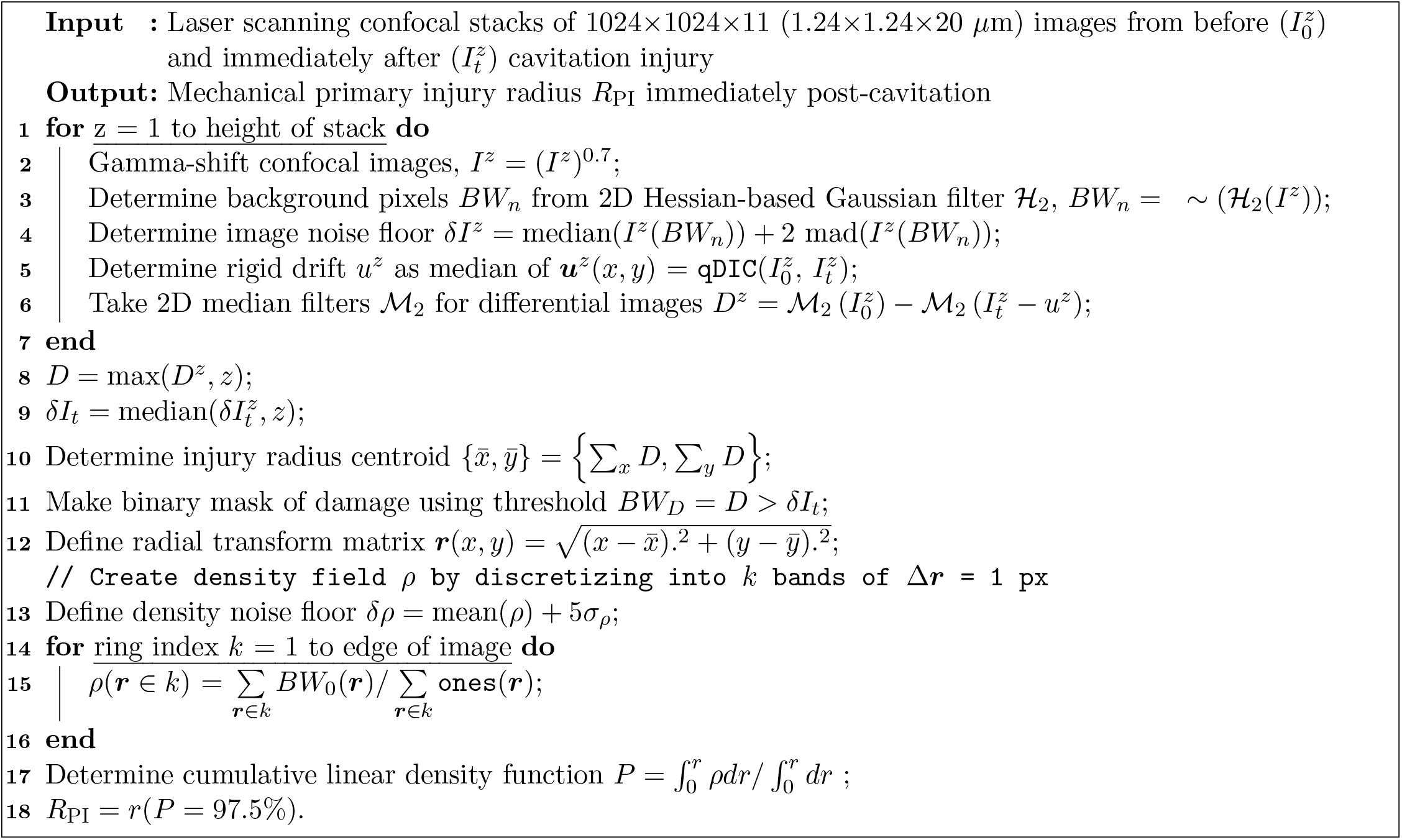
Calculating*R*_PI_ from laser scanning confocal image stacks.

### 3D Image processing

The general processing procedure for determining the primary injury radius from the reference and post-injury confocal image stacks of cells stained with Calcein AM is outlined in Algorithm 2.

Each z-image in an image stack was first gamma-shifted by a factor of 0.7, and subsequently processed by a Hessian-based Gaussian filter [66] kernel with *σ* ranging from 1 to 3 to determine a slice-wise noise floor. Due in part to drift, switching imaging optics, and to the cavitation event itself, some rigid translation occurred between each image before and after cavitation; registering image drift is a common part of image differencing techniques to get accurate results [67]. For this study, we corrected for rigid motion drift by running a quality-factor based digital image correlation method [28] and shifting the second image by the determined median image displacement. Corresponding images in the reference configuration were then compared to rigid-motion-corrected images in the damaged configuration at times immediately after cavitation. Differences between the image pairs, called *differential* images, were then median-filtered, summed in the stack-direction, and thresholded automatically by the Hessian-determined noise floor. A binary mask was constructed as the values in the differential image stack sum, *D*, above the threshold. This binary mask represents all pixels that significantly drop in intensity between the undamaged and damaged images, and physically represents local ceasing of intracellular esterase activity. To assess disruption of esterase activity radially, the epicenter of the cavitation injury 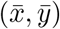 is calculated from the 2D median of the cumulative sum of the rows and columns of ***D***.

**Algorithm 3:**
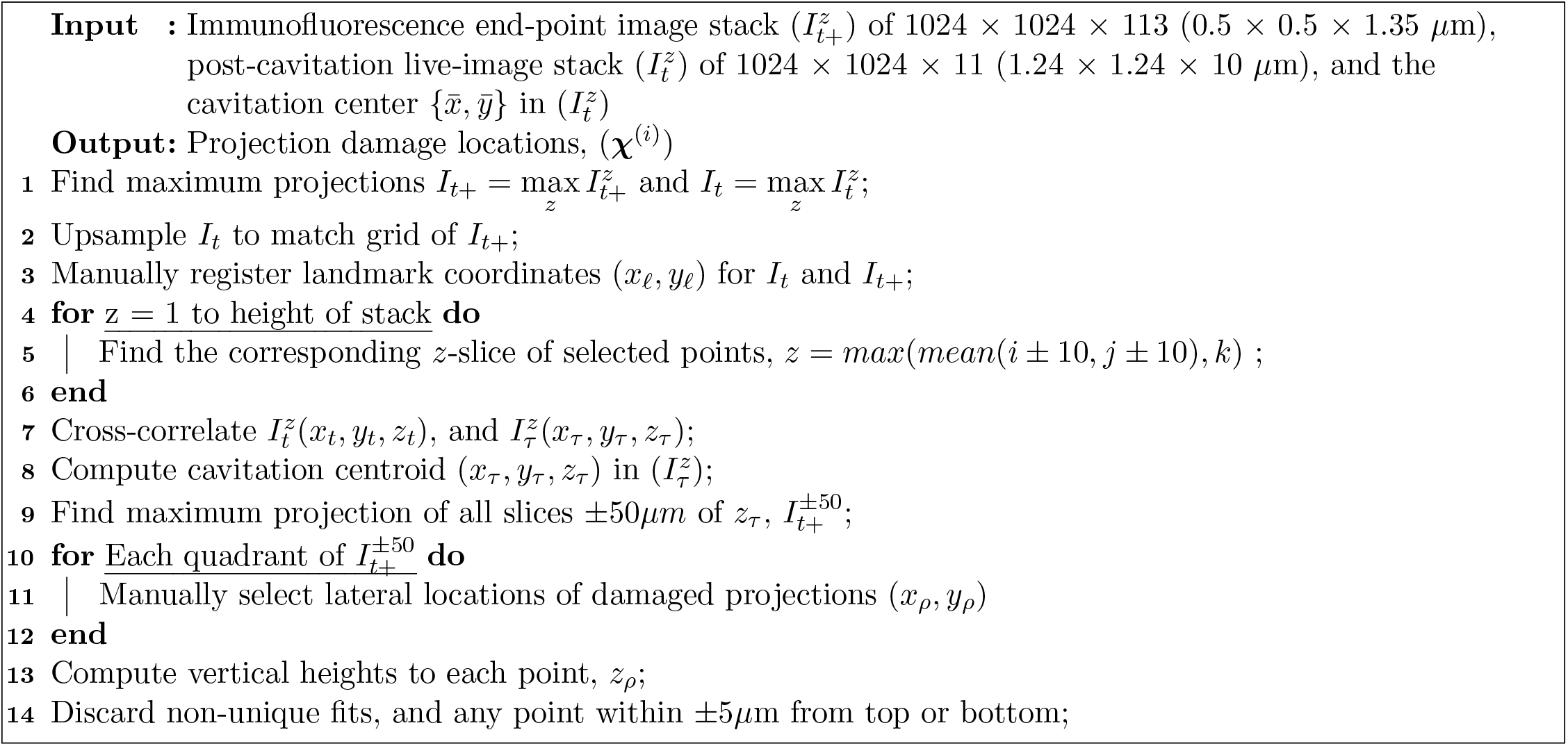
Calculating radial positions of damaged neural cellular projections.

A distance transform matrix ***r***(*x, y*) is constructed as the distance away from the bubble epicenter location for each pixel (*x, y*) in *D*. From the center of the bubble at *r* = 0, it is convenient to take the relative density difference as a radial measure of damage, that is, the high density of fluorescent intensity loss concentrated in the bubble region is considered the damaged zone. Intensity loss density *ρ* was determined by defining Δ***r***-width concentric rings, summing the lost pixels (where the mask is 1), and normalizing by the number of pixels in each ring. The density noise floor is user-adjustable, but was conservatively taken as 5 standard deviations above the median far-field density in this study. The radius of mechanical disruption or primary injury of the plasma membrane immediately after cavitation, *R*_PI_, was then defined via the linear integral of the radial density; specifically, where 97.5% of the cumulative fluorescent intensity loss (i.e. two standard deviations above the median) occurs.

The general immunofluorescence image processing procedure for determining the locations of damaged projections is outlined in Algorithm 3. Immunofluorescence end-point image stacks 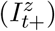 of 1024 × 1024 × 113 (0.5 × 0.5 × 1.35 *μ*m) are used to manually determine the location in three dimensional space of damaged projections for each immunofluorescence marker. Here, a damaged projection is defined as any continuous projection originating from a cell soma that terminates abrubtly and without intuitive cause to the observer with experience examining the healthy morphological state of 3D neural cell cultures. A calcein AM post-cavitation live-image stack 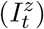 of 1024 × 1024 × 11 (1.24 × 1.24 × 10 *μ*m) is upsampled to match the grid size of 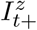 and manually registered with a combination of manual landmark matching and image cross-correlation. An input cavitation epicenter 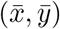 — assumed to fall in the mid-plane of 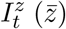—is then converted from the known coordinates in 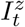 to the corresponding coordinates in 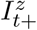. Maximum projections are computed from all slices of 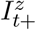 determined to fall within ±50*μ*m of 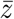 and a quadrant-wise manual point selection is completed to localize the lateral location of damaged projections. All lateral(*x* − *y*) positions manually identified are registered in along the *z*-axis by computing the maximum value of the median intensity within a window constructed about each *x, y* point along *z*. Any point determined to fall within the top or bottom 5*μ*m of the image stack are neglected to avoid image clipping; additionally, any point without a unique maximum value is excluded from further processing.

## Funding

Office of Naval Research grant N00014-16-1-2872 (JBE, HCC, CF)

Office of Naval Research grant N00014-17-1-2058 (SB, CF)

Office of Naval Research grant N00014-18-1-2630 (HCC, CF)

Office of Naval Research grant N00014-16-1-2869 (MTS, CF)

Office of Naval Research grant N00014-17-1-2644 (MTS, CF)

Graduate Assistance in Areas of National Need (GAANN) fellowship, Brown University Institute for Molecular and Nanoscale Innovation (JBE)

## Author contributions

Conceptualization: JBE, HCC, MTS, CF

Methodology: JBE, HCC, MTS, CF

Investigation: JBE, HCC, MTS, SB

Visualization: JBE, HCC

Supervision: JBE, CF

Writing—original draft: JBE, HCC, CF

Writing—review & editing: JBE, HCC, MTS, SB, CF

## Competing interests

Authors declare that they have no competing interests.

## Data and materials availability

All data and micrographs are available through the University of Wisconsin Digital Collections Center, University of Wisconsin System (UW@MINDS) dataset entitled “Neural cell injury pathology due to high-rate mechanical loading” at doi.org/10.21231/z3c6-yy40.

## Notes

### Competing Interest Statement

The authors have declared no competing interest.

### Summary of Updates

Minor abstract edit.

## References

[1] J. D. Humphrey, E. R. Dufresne, M. A. Schwartz, Mechanotransduction and extracellular matrix homeostasis. Nat. Rev. Mol. Cell Biol. 15(12), 802–812 (2014).

[2] S. J. Tan, A. C. Chang, S. M. Anderson, C. M. Miller, L. S. Prahl, D. J. Odde, A. R. Dunn, Regulation and dynamics of force transmission at individual cell-matrix adhesion bonds. Sci. Adv. 6(20), eaax0317 (2020).

[3] S. Okuda, N. Takata, Y. Hasegawa, M. Kawada, Y. Inoue, T. Adachi, Y. Sasai, M. Eiraku, Strain-triggered mechanical feedback in self-organizing optic-cup morphogenesis. Sci. Adv. 4(11):eaau1354 (2018).

[4] L. J. Simpson, J. S. Reader, E. Tzima, Mechanical Regulation of Protein Translation in the Cardiovascular System. Front. Cell Dev. Biol. 8(34) (2020).

[5] O. Chaudhuri, S. T. Koshy, C. Branco Da Cunha, J. Shin, C. S. Verbeke, K. H. Allison, D. J. Mooney Extracellular matrix stiffness and composition jointly regulate the induction of malignant phenotypes in mammary epithelium. Nat Mat 13, 970–978 (2014).

[6] E. Bar-Kochba, M. T. Scimone, J. B. Estrada, C. Franck Strain and rate-dependent neuronal injury in a 3D in vitro compression model of traumatic brain injury. Sci Rep 6, 30550 (2016).

[7] M. T. Scimone, H. C. Cramer III, E. Bar-Kochba, R. Amezcua, J. B. Estrada, C. Franck, Modular approach for resolving and mapping complex neural and other cellular structures and their associated deformation fields in three dimensions. Nat. Protoc. 13(12), 3042–3064 (2018).

[8] C. E. Brennen, Cavitation in Medicine. Interface Focus. 5, 20150022 (2015).

[9] D. L. Miller, R. M. Thomas, Ultrasound contrast agents nucleate inertial cavitation in vitro. Ultrasound Med. Biol. 21(8), 1059–1065 (1995).

[10] M. W. Miller, D. L. Miller, A. A. Brayman, A review of in vitro bioeffects of inertial ultrasonic cavitation from a mechanistic perspective. Ultrasound Med Biol 22(9), 1131–1154 (1996).

[11] C. C. Church, Spontaneous homogeneous nucleation, inertial cavitation and the safety of diagnostic ultrasound. Ultrasound Med Biol 28(10), 1349–1364 (2002).

[12] M. Trendowski, The promise of sonodynamic therapy. Cancer Metastasis Rev. 33, 143–160 (2014).

[13] C. C. Church, A theoretical study of cavitation generated by an extracorporeal shock wave lithotripter. J. Acoust. Soc. Am. 86(1), 215–227 (1989).

[14] Z. Xu, Z. Fan, T. L. Hall, F. Winterroth, J. B. Fowlkes, C. A. Cain, Size measurement of tissue debris particles generated from pulsed ultrasound cavitational therapy – histotripsy. Ultrasound Med. Biol. 35(2), 245–255 (2009).

[15] A. D. Maxwell, C. A. Cain, A. P. Duryea, L. Yuan, H. S. Gurm, Z. Xu, Noninvasive thrombolysis using pulsed ultrasound cavitation therapy – histotripsy. Ultrasound in Med. and Biol. 35(12), 1982–1994 (2009).

[16] V. Venugopalan, A. Guerra III, K. Nahen, A. Vogel, Role of laser-induced plasma formation in pulsed cellular microsurgery and micromanipulation. Phys. Rev. Lett. 88(7), 078103 (2002).

[17] A. V. Cherian, K. R. Rau, Pulsed-laser-induced damage in rat corneas: time-resolved imaging of physical effects and acute biological response. J. Biomed. Opt. 13(2), 024009 (2008).

[18] J. R. Sukovich, C. A. Cain, A. S. Pandey, N. Chaudhary, S. Camelo-Piragua, S. P. Allen, T. L. Hall, J. Snell, Z. Xu, J. M. Cannata, D. Teofilovic, Bertolina JA, Kassell N, and Xu Z(2018). In vivo histotrypsy brain treatment. J Neurosurg.: 1–8.

[19] J. J. Macoskey, S. W. Choi, T. L. Hall, E. Vlaisavljevich, J. E. Lundt, F. T. Lee Jr, E. Johnsen, C. A. Cain, Z. Xu, Using the cavitation collapse time to indicate extent of histotripsy-induced tissue fractionation. Phys. Med. Biol. 63(5): 55013 (2019).

[20] S. B. Shively, I. Horkayne-Szakaly, R. B. Jones, J. P. Kelly, R. C. Armstrong, D. P. Perl, Characteristic of interface astroglial scarring in the human brain after blast exposure: a post-mortem case series.Lancet Neurol. 15(9):944–953 (2016).

[21] A. Dagro, J. Wilkerson, A computational investigation of strain concentration in the brain in response to a rapid temperature rise. J. Mech. Behav. Biomed. 115, 104228 (2021).

[22] M. Nyein, A. M. Jason, L. Yu, C. M. Pita, J. D. Joannopoulos, D. F. Moore, R. A. Radovitzky, In silico investigation of intracranial blast mitigation with relevance to military traumatic brain injury. Proc. Natl. Acad. Sci. U.S.A. 107, 20703–20708 (2010).

[23] M. C. Rice, E. M. Arruda, M. D. Thouless, The use of visco-elastic materials for the design of helmets and packaging, J Mech Phys Solids 141, 103966 (2020).

[24] J. B. Estrada, C. Barajas, D. L. Henann, E. Johnsen, Franck C, High strain-rate soft material characterization via inertial cavitation. J. Mech. Phys. Solids 112, 291–317 (2018).

[25] M. T. Scimone, H. C. Cramer III, P. Hopkins, J. B. Estrada, C. Franck, Application of mild hypothermia successfully mitigates neural injury in a 3D in vitro model of traumatic brain injury. PLoS ONE 15(4): e0229520 (2020).

[26] J. B. Keller, M. Miksis, Bubble oscillations of large amplitude. J. Soc. Acoust. Amer. 68(2), 628–633 (1980).

[27] A. Prosperetti, A. Lezzi, Bubble dynamics in a compressible liquid. Part 1. First-order theory. J. Fluid Mech. 168, 457–478 (1986).

[28] A. K. Landauer, M. Patel, D. L. Henann, C. Franck, A q-factor-based digital image correlation algorithm (qDIC) for resolving finite deformations with degenerate speckle patterns. Exp. Mech. 58(5), 815–830 (2018).

[29] E. Gurdjian, H. Lissner, R. Latimer, B. Haddad, J. Webster, Quantitative determination of acceleration and intercranial pressure in experimental head injury. Neurology 3, 417–423 (1953).

[30] B. Morrison III, B. S. Elkin, J. P. Dollé, M. L. Yarmush, In vitro models of traumatic brain injury. Annu. Rev. Biomed. Eng. 3, 91–126 (2011).

[31] M. C. LaPlaca, D. K. Cullen, J. J. McLoughlin, R. S. Cargill, high-rate shear strain of three-dimensional neural cell cultures: a new in vitro traumatic brain injury model. J. Biomech. 38, 1093–105 (2005).

[32] D. K. Cullen, C. M. Simon, M. C. LaPlaca, Strain rate-dependent induction of reactive astrogliosis and cell death in three-dimensional neuronal–astrocytic co-cultures. Brain Research 1158, 103–115 (2007).

[33] S. E. Legget, M. Patel, T. M. Valentin, L. Gamboa, A. S. Khoo, E. K. Williams, C. Franck, I. Y. Wong, Mechanophenotyping of 3D multicellular clusters using displacement arrays of rendered tractions. Proc. Natl. Acad. Sci. U.S.A. 117(11) 5655–5663 (2020).

[34] R. M. Wright, K. T. Ramesh, An axonal strain injury criterion for traumatic brain injury. Biomech. Model Mechanobiol. 11(1-2), 245–260 (2012).

[35] R. W. Carlsen, A. L. Fawzi, A. Wan, H. Kesari, C. Franck, A quantitative relatioship between rotational head kinematics and brain tissue strain from 2D parametric finite element analysis. Brain Mulitiphysics 2: 100024 (2021).

[36] M. B. Panzer, B. S. Myers, B. P. Capehart, C. R. Bass, Development of a Finite Element Model for Blast Brain Injury and the Effects of CSF Cavitation. Ann. Biomed. Eng. 40, 1530–1544 (2012).

[37] D. H. Smith, D. F. Meaney, W. H. Shull, Diffuse Axonal Injury in Head Trauma. Journal of Head Trauma Rehabilitation 18 (4), 307–316 (2003).

[38] Y. T. Wu, A. Adnan, Damage and failure of axonal microtubule under extreme high strain rate: an in-silico molecular dynamics study. Sci. Rep. 8, 12260 (2018).

[39] I. K. Khan, S. F. Ferdous, A. Adnan, Mechanical Behavior of actin and spectrin subjected to high strain rate: A molecular dynamics simulation study. Comput. Struct. Biotechnol. J. 19, 1738–1749 (2021).

[40] M. I. Khan, F. Hasan, K. A. H. Al Mahmud, A. Adnan, Recent Computational approaches on mechanical behavior of axonal cytoskeletal components of neuron: a brief review. Multiscale Sci. Eng. 2, 199–213 (2020).

[41] D. M. Geddes, R. S. Cargill II, M. C. LaPlaca, high-rate shear strain of three-dimensional neural cell cultures: a new in vitro traumatic brain injury model. J. Biomech. 38(5), 1093–1105 (2003).

[42] B. Morrison III, H. L. Cater, C. C. B. Wand, F. C. Thomas, C. T. Hung, G. A. Ateshian, L. E. Sundstrom, A Tissue Level Tolerance Criterion for Living Brain Developed with an In Vitro Model of Traumatic Mechanical Loading. Stapp Car Crash Journal 47, 93–105 (2003).

[43] D. Ambrosi, G. A. Ateshian, E. M. Arruda, S. C. Cowin, J. Dumais, A. Goriely, G. A. Holzapfel, J. D. Humphrey, E. Kuhl, J. E. Olberding, L. A. Taber, K. Garikipati K, Perspectives on biological growth and remodeling J. Mech. Phys. Solids. 59(4), 863–883 (2011).

[44] S. Budday, R. Nay, R. de Rooij, P. Steinmann, T. Wyrobek, T. C. Ovaert, E. Kuhl, Mechanical properties of gray and white matter brain tissue by indentation. J Mech Behav Biomed Mater 46, 318–330 (2015).

[45] E. Spedden, J. D. White, E. N. Naumova, D. L. Kaplan, C. Staii, Elasticity maps of living neurons measured by combined Fluorescence and atomic force microscopy. Biophys. J. 103(5), 868–877 (2012).

[46] H. Ouyang, E. Nauman, R. Shi, Contribution of cytoskeletal elements to the axonal mechanical properties. J. Biol. Eng. 7:21 (2013).

[47] R. Bernal, P. A. Pullarkat, F. Melo, Mechanical properties of axons. Phys. Rev. Lett. 99, 018301 (2007).

[48] S. Nam, K. H. Hu, M. J. Butte, O. Chaudhuri, Strain-enhanced stress relaxation impacts nonlinear elasticity in collagen gels. Proc. Natl. Acad. Sci. U.S.A. 113(20), 5492–5497 (2015).

[49] D. Velegol, F. Lanni, Cell traction forces on soft biomaterials. I. Microrheology of type I collagen gels. Biophys. J. 81(3), 1786–1792 (2001).

[50] G. Lai, Y. Li, G. Li, Effect of concentration and temperature on the rheological behavior of collagen solutionInt. J. Biol. Macromol. 42(3), 285–291 (2008).

[51] S. A. A. Machado, V. C. A. Martins, A. M. G. Plepis, Thermal and Rheological behavior of collagen and chitosan blends. J. Therm. Anal. Cal. 67, 1418–2874 (2002).

[52] Y. B. Lu, K. Franze, G. Seifert, C. Steinhauser, F. Kirchhoff, H. Wolburg, J. Guck, P. Janmey, E. Q. Wei, J. Kas, A. Reichenbach, Viscoelastic properties of individual glial cells and neurons in the CNS. Proc. Natl. Acad. Sci. U.S.A. 103(47) 17759–17764 (2006).

[53] K. Xu, G. Zhong, X. Zhuang, Actin, Spectrin, and associated proteins form a periodic cytoskeletal structure in axons. Science 339(6118), 452–456 (2013).

[54] I. Peng, L. I. Binder, M. M. Black, Biochemical and immunological analyses of cytoskeletal domains of neurons. J. Cell Biol. 102(1), 252–262 (1986).

[55] L. C. Kapitein, C. C. Hoogenraad, Building the neuronal microtubule cytoskeleton. Neuron 87(3), 492–506 (2015).

[56] A. Koleske, Molecular mechanisms of dendrite stability Nat. Reviews 14, 536–550 (2013).

[57] H. Ahmadzadeh, D. H. Smith, V. B. Shenoy, Viscoelasticity of tau proteins leads to strain rate-dependent breaking of microtubules during axonal stretch injury: predictions from a mathematical model. Biophys. J. 106(5), 1123–1133 (2014).

[58] L. Mancia, E. Vlaisavljevich, N. Yousefi, M. Rodriguez, T. J. Ziemlewicz, F. T. Lee, D Hennan, C. Franck, Z. Xu, E. Johnsen, Modeling tissue-selective cavitation damage.Phys. Med. & Biol. 64(22), 2250001 (2019).

[59] D. F. Meaney, B. Morrison III, C. D. Bass, The mechanics of traumatic brain injury: a review of what we know and what we need to know for reducing its societal burden. J Biomech. Eng. 136(2), 0210081–02100814 (2014).

[60] X. Nie, B. SanBorn, T. Weerasooriya, W. Chen, High-rate bulk and shear responses of bovine brain tissue. Int. J. Impact Eng. 53, 56–61 (2013).

[61] Z. Wang, J. B. Estrada, E. M. Arruda, K. Garikipati, Inference of deformation mechanisms and constitutive response of soft material surrogates of biological tissue by full-field characterization and data-driven variational system identification. J. Mech. Phys. Solids 153, 104474 (2021).

[62] M. C. LaPlaca, V. N. Vernekar, J. T. Shoemaker, D. K. Cullen, Three-dimensional neuronal cultures. Methods in bioengineering: 3D tissue engineering. Artech House, Norwood, MA, 187–204 (2010).

[63] Y-T. Dingle, M. E. Boutin, A. M. Chirila, L. L. Livi, N. R. Labriola, L. M. Jakubek, J. R. Morgan, E. M. Darling, J. A. Kauer, D. Hoffman-Kim, Three-dimensional neuronal spheroid culture: an in vitro model for cortical studies. Tissue Eng. Part C Methods 21(12), 1275–1283 (2015).

[64] G. Taubin, Estimation of planar curves, surfaces, and nonplanar space curves defined by implicit equations with applications to edge and range image segmentation. IEEE Trans. Pattern Anal. Mach. Intell. 13(11), 1115–1138 (1991).

[65] J. Yang, H. C. Cramer III, C. Franck, Extracting non-linear viscoelastic material properties from violently collapsing cavitation bubbles. Extreme Mech. Lett. 39, 100839 (2020).

[66] A. F. Frangi, W. J. Niessen, K. L. Vincken, M. A. Viergever, Multiscale vessel enhancement filtering. In: Wells W.M., Colchester A., Delp S. (eds) Medical Image Computing and Computer-Assisted Intervention — MICCAI’98 MICCAI, (2006).

[67] R. J. Radke, A. Srinivas, O. Al-Kofahni, B. Roysam, Image change detection algorithms: a systematic survey. IEEE Trans. Image Process 14(3), 294–307 (2005).

